# Interconversion of tripartite and bipartite systems for site-specific *O*-sialylation of flagellin in Gram-negative and Gram-positive bacteria

**DOI:** 10.1101/2023.12.03.569772

**Authors:** Jovelyn Unay, Nicolas Kint, Patrick Viollier

## Abstract

Many bacteria decorate flagellin with sialic acid-like sugars such as pseudaminic acid (Pse) by O-glycosylation on serine or threonine residues. Evidence for sufficiency of sialylation by a conserved flagellin glycosyltransferase (fGT) system is lacking, presumably because of (a) missing component(s). Here, we reconstituted two Maf-type fGTs from the Gram-negative bacterium *Shewanella oneidensis* MR-1 in a heterologous host producing a Pse donor sugar. While Maf-1 is sufficient for flagellin glycosylation, Maf-2 reconstitution requires a newly identified, *cis*-encoded and conserved specificity factor GlfM, predicted to form a four-helix bundle. While GlfM binds Maf-2 to form a ternary complex with flagellin, the C-terminal tetratricopeptide repeat (TPR) domain of Maf-1 confers flagellin acceptor and O-glycosylation specificity at preferred serine residues. GlfM from Gram-negative and Gram-positive bacteria are functional, providing evidence for convergent evolution of specialized flagellin modification systems with acceptor serine selectivity, while also shaping the interconversion of bacterial tripartite and bipartite O-glycosylation systems.

## INTRODUCTION

Covalent post-translational modification of proteins not only alters chemical properties of proteins, but also frequently their function. In its regulatory capacity, post-translational modification of proteins is designed to impose changes in activity on selected proteins only. In turn, this necessitates the selective recognition of the protein substrates as acceptors of such covalent modification to prevent the inappropriate modification of other proteins.

Various forms of post-translational modifications have been described for eukaryotic and prokaryotic cells, yet protein glycosylation in bacterial cells remains understudied despite its frequent occurrence^1,2^. One common type of protein glycosylation occurs on flagellins, the subunit of the flagellar filament, by O-linked glycosylation at specific acceptor serine or threonine residues. In its most common form, flagellin glycosylation differs fundamentally to other bacterial protein glycosylation systems in two ways. First, the donor glycan that is covalently coupled to flagellin is a derivative of sialic acid, a class of nine-carbon (α-keto) acidic sugars featuring acetamido linkages^1,3^. Second, glycosyl transfer occurs in the bacterial cytoplasm, before flagellin is secreted and assembled into the surface-exposed flagellar filament^4–6^.

The flagellar assembly machinery is an envelope-spanning structure that houses a protein translocation system and rotary engine embedded in the inner membrane, as well as the drive shaft and bushings that traverse the cell wall (peptidoglycan, PG, layer) and the outer membrane in the case of Gram-negative (diderm) bacteria, collectively referred to as the basal body ^7–10^. The surface exposed hook functions as a flexible joint that transmits torque from the rotor to the long flagellar filament that acts as a propeller. For filament assembly to occur, dedicated secretion chaperones direct flagellin subunits through the hollow conduit in the hook-basal-body (HBB) traversing the PG layer where they are received by specialized polymerization chaperones^11,12^, also known as capping proteins, located at the tip of the nascent filament. These polymerization chaperones catalyse the sequential incorporation of flagellin at the distal tip, forming a tubular filament composed of 11 protofilaments adopting different superhelical states^13^.

Recent cryo-electron microscopic structural studies aimed at defining these states of the flagellar filament unexpectedly revealed flagellin subunits featuring post-translational modifications manifested as extra electron densities, thought to reflect covalently coupled glycans ^14,15^. Indeed, flagellar filaments are post-translationally modified by sialic acid derivatives called pseudaminic acid (Pse, 5,7-diacetamido-3,5,7,9-tetradeoxy-L-glycero-L-manno-nonulosonic acid) or its stereoisomer legionaminic acid (Leg)^1,3,5^. Illumination of the mysterious mechanism by which flagellins are specifically selected from other cytoplasmic proteins for covalent modification with Pse or Leg awaited the discovery and reconstitution of glycosylation by the FlmG-class of flagellin glycosyltransferases (fGTs) encoded in the α- proteobacteria *Caulobacter crescentus* and *Brevundimonas subvibroides*^4,16^. FlmGs feature an overt and genetically separable two-domain architecture: an N-terminal tetratricopeptide repeat (TPR) that mediates flagellin binding, while a C-terminal glycosyltransferase (GT) domain adopting a GT-B fold executes the glycosylation reaction with preference for Pse or Leg^5,16,17^. Importantly, FlmG can glycosylate flagellin when expressed in heterologous hosts synthesizing a suitable donor glycan: a Pse derivative in the case of *C. crescentus* FlmG and a Leg derivative for *B. subvibroides* FlmG. Since FlmGs are cytoplasmic protein GTs that use soluble sialic acid as donor glycan, these glycosylation systems differ fundamentally in topology from the O- or N-glycosyl transferases executed by periplasmic protein GTs that depend on donor glycan that is membrane anchored^5,18^.

While the distribution of FlmG enzymes across bacterial genomes is rather narrow and confined to the *Caulobacterales* order of the α-proteobacteria, many other bacterial species also glycosylate flagellins. Their flagellar gene clusters instead encode a putative fGT called the <u>m</u>otility <u>a</u>ssociated factor (Maf) harbouring a GT-A type fold, also known as the MAF_flag10 domain or <u>D</u>omain of <u>U</u>nknown <u>F</u>unction (DUF) 115 ^5,17,19–21^. Some bacteria encode multiple Mafs in the same flagellar gene cluster ^18,19,22,23^ that have been shown to control flagellation, likely by controlling flagellin abundance and/or modification^19,20,23–25^. Inactivation of Pse or Leg biosynthesis pathways can also lead to flagellation and flagellin glycosylation defects, suggesting that Mafs glycosylate flagellin with Pse or Leg ^19,20,22,23,26–29^. Mafs are widely distributed among Gram-negative lineages including certain orders of α- proteobacteria (e.g., *Magnetospirillum magneticum*), ε-proteobacteria (e.g., *Campylobacter jejuni*) and γ-proteobacteria (e.g., *Shewanella oneidensis*), but also in certain spirochetes for example *Treponema denticola*. Maf orthologs are also encoded in the flagellar gene clusters of Gram-positive bacteria belonging to the genera *Clostridia*, *Kurthia* or *Geobacillus*.

How Mafs function has remained largely obscure, in part because the determinants for the acceptor (flagellin) and the donor (Pse or Leg) are unknown and because it is unknown whether additional components are required for efficient reconstitution of flagellin glycosylation by Maf with a suitable Leg or Pse donor. Confounding the dissection of Maf specificity is the multiplicity of Maf enzymes encoded in certain genomes, for example *S. oneidensis* MR-1, but also the diversity of Pse and Leg derivatives that are produced by different bacteria and the fact that some bacteria produce both Pse and Leg. In some cases, bacteria not only use Pse or Leg to decorate flagellin (also known as H -antigen), but also to modify other cell surface structures such as capsules (the K-antigen) or the O-antigen component of lipopolysaccharide (LPS)^1,3^.

Here, we report the dissection and reconstitution of Pse-dependent flagellin glycosylation by Maf-1 and Maf-2 from *S. oneidensis* MR-1. We show that Maf-1 and Maf-2 both glycosylate flagellin with a Pse derivative, even in a heterologous system. However, they target distinct serine residues through different flagellin binding domains. We find that Maf-1 functions as a bipartite protein glycosylation system that interacts directly with flagellin, while Maf-2 constitutes a tripartite system that can only be functionally reconstituted in the presence of a previously unknown specificity factor, GlfM, that promotes complex formation between Maf-2 and flagellin. The coding sequences for GlfM and Maf-2 are juxtaposed in the genomes of Gram-negative and Gram-positive bacteria, pointing to an evolutionary interconversion between bipartite and tripartite glycosylation systems with an optional control element that bipartite systems lack.

## RESULTS

### Pse-dependent flagellin glycosylation by *S. oneidensis* Maf-1, but not by Maf-2 orthologs

A simple and straightforward strategy to assess the functionality and sufficiency of flagellin glycosyltransferases is to co-express them with their cognate flagellin in a heterologous host producing a suitable donor glycan. *S. oneidensis* MR-1 is an intriguing system for flagellin glycosylation because it encodes two Maf paralogs, both expressed from within the flagellar gene cluster (**Fig. 1A**) and flanking Pse biosynthesis genes. Since this organization suggests that Pse could serve as glycosyl donor for the Maf, we co-expressed each Maf with the corresponding (FLAG-tagged) flagellin in *C. crescentus* wild-type (*WT*) cells that produce Pse or Δ*neuB* mutant cells that lack Pse^4^. The migration of flagellin on SDS-PAGE was probed by immunoblotting using monoclonal antibodies to the FLAG-epitope. We noted a (small) retardation of flagellin migration when co-expressed with Maf-1 in *WT* cells *versus* Δ*neuB* cells (**Supplementary Fig. S1A**). Moreover, FLAG-tagged flagellin also migrates slower when *S. oneidensis* Maf-1 is expressed in an *E. coli* host containing a synthetic operon encoding the six Pse-biosynthesis genes from *C. crescentus*^4^ (**Supplementary Fig. S1B**). Surprisingly, however, we could not detect any Pse-dependent change in flagellin migration when Maf-2 is co-expressed with flagellin in *C. crescentus WT* cells *versus* Δ*neuB* cells (**Supplementary Fig. S1C**). We conclude that *S. oneidensis* Maf-1 is active and sufficient for flagellin glycosylation in *C. crescentus*, whereas Maf-2 is either inactive or requires a helper protein.

**Figure 1.**
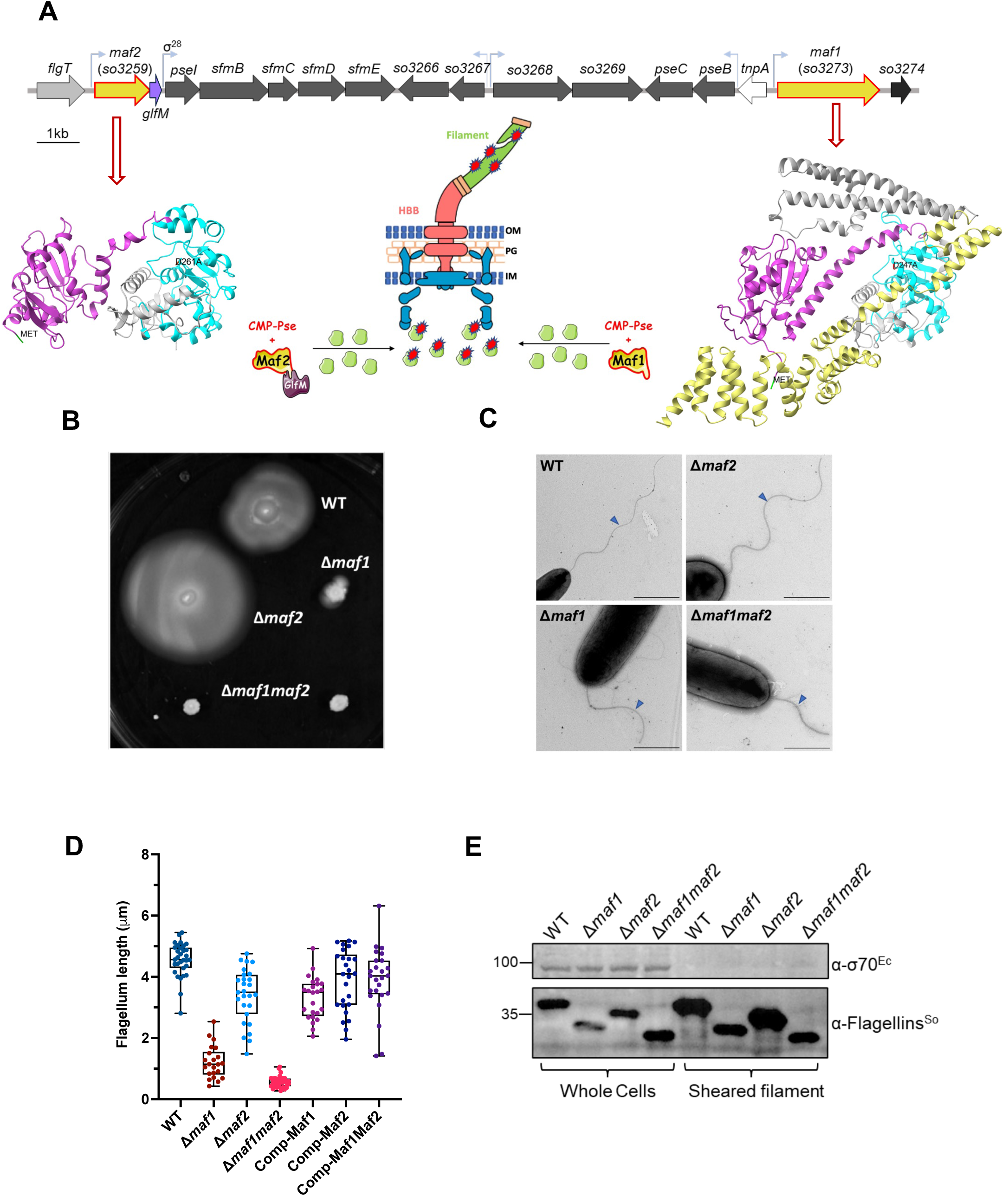
Maf-1 and Maf-2 affect motility and flagellin modification in *S. oneidensis* MR- 1. **A)** Genetic map of *S. oneidensis* MR-1 showing the location of the genes encoding Maf- 1 (*So3273*) and Maf-2 (*So3259*) depicted in yellow. Genes *maf-1* and *maf-2* are within the large genetic motility island, flank the putative pseudaminic acid (Pse) biosynthesis gene cluster of *S. oneidensis* MR-1 (also called as *<u>S</u>hewanella* flagellin <u>m</u>odification or *sfm* operon, ^22^). Putative Pse biosynthesis genes are depicted in dark grey while the gene encoding GlfM is coloured in purple. Structures of Maf-1 and Maf-2 are predicted using AlphaFold v2. The N-terminal region of both proteins is coloured in magenta while the signature MAF-Flag10 domain or <u>D</u>omain of <u>Un</u>known <u>F</u>unction (DUF) 115 is shown in turquoise and the predicted tetratrico<u>p</u>eptide repeat (TPR) domain of Maf1 is coloured in yellow. The figure also depicts the glycosylation of flagellin by either Maf- 1 or Maf-2-GlfM before export via the flagellar secretion apparatus, through the hook basal body (HBB) and along the filament channel, followed by its polymerization at the distal tip of the flagellar filament by a capping protein. **B)** Motility assay of *S. oneidensis WT* and mutant strains on swarm agar (0.25% LB agar). Strains were allowed to grow overnight at 30°C followed by overnight incubation at RT. This assay is a representative of at least three independent experiments. **C)** Flagellar filaments of *S. oneidensis WT* and *maf* mutants visualized by transmission electron microscopy. Filaments are indicated with blue caret. Scale bars are 1μm. **D)** Graph showing flagellar filament length measurement in *S. oneidensis WT*, *maf* mutants and complemented strains. Filament length was measured in 22-32 cells per strain using Fiji^55^. **E)** Immunoblot of flagellins in extracts from *WT* and mutant cells and in the supernatant following their release after mechanically shearing flagellar filaments from the cell surface. Blots were probed with polyclonal antibodies to *S. oneidensis* FlaA/B (α- Flagellins^So^)^22^. As control for the shearing, blots were also probed with a monoclonal clonal antibody to σ^70^ of *E. coli* (α-σ^70Ec^) to determine whether shearing using a syringe- needle resulted in cells lysis, thereby liberating the cytoplasmic content including σ^70^.

Interestingly, the predicted domain structures of Maf-2 and Maf-1 differ. While both Mafs harbour the signature motility associated factor, MAF_flag10 domain or DUF 115 predicted to have a GT-A type glycosyltransferase-like fold (**Fig. 1A**), they are only 21% similar (**Supplementary Fig. S2**). Moreover, Maf-1 is 822 residues long, while Maf-2 is a 441-residue protein. HHpred analyses suggests that Maf1 harbours a C-terminal tetratrico<u>p</u>eptide repeat (TPR) domain (550-820 aa), while Maf-2 has no C-terminal domain after the signature MAF_flag10 domain. Since a TPR domain was recently shown to be critical and sufficient for flagellin binding by the less conserved FlmG-class of fGTs discovered in *C. crescentus* ^4^ and *B. subvibrioides* ^16^, we speculated that the TPR could represent as the flagellin binding determinant in Maf-1, while there is no apparent flagellin binding determinant on the polypeptide of Maf-2.

### Maf-1 and Maf-2 have opposing roles on motility in *S. oneidensis* MR-1

To explore whether Maf-2 contributes to flagellin glycosylation *in S. oneidensis* MR-1, we constructed *S. oneidensis* MR-1 mutant strains with single and double deletions (Δ*maf-1*, Δ*maf-2* and Δ*maf-1*Δ*maf-2*) and probed for defects in flagellum-mediated swarming of these mutants on semi-solid (motility or swarm) agar. We observed that the Δ*maf-1* and Δ*maf- 1*Δ*maf-2* strains form a compact cluster of cells (**Fig. 1B**). By contrast, the expanded swarms of *WT* and Δ*maf-2* single mutant strain indicate proper flagellar function. Interestingly, the Δ*maf-2* swarms appeared slightly larger than the *WT* swarms, suggesting that Maf-2 can have negative impact on motility. To visualize the flagella on *WT* and mutant cells we conducted transmission electron microscopy (TEM) and observed substantially shortened flagellar filaments (ca 1-2μm) on Δ*maf-1* cells compared to the long filament on *WT* cells (ca 2-6 μm; **Fig. 1C, 1D**). The flagellar filament of Δ*maf-1*Δ*maf-2* cells is even shorter than that of Δ*maf-1* cells (**Fig. 1D**), but the filaments Δ*maf2* cells (2-4μm) is hardly shorter to that of *WT* cells (**Fig. 1C, +D**). Upon introducing the corresponding complementing plasmids, all *maf* mutants feature a flagellar filament length comparable to *WT* cells (**Fig. 1D**).

To correlate these flagellar filament lengths with flagellin levels, we sheared flagellar filaments from *WT* and mutant cells using a syringe needle and probed the supernatant for flagellin abundance by immunoblotting using polyclonal antibodies to the flagellins FlaA/B of *S. oneidensis* MR-1 ^22^. As control we also separated extracts from *WT* and mutant cells by SDS-PAGE and probed for corresponding changes in migration and abundance of cell associated flagellin on immunoblotting. As seen in **Fig. 1E**, flagellins are clearly less abundant in Δ*maf1* and Δ*maf-1*Δ*maf-2* cells and in the supernatant of sheared cells. Importantly, these experiments also revealed the increased migration of flagellins from Δ*maf1* and Δ*maf-1*Δ*maf- 2* cells compared to the flagellin of *WT* cells. Moreover, the flagellins of Δ*maf-2* cells also migrate faster than the flagellins of *WT* cells and the flagellins of Δ*maf-1*Δ*maf-2* cells migrate even faster than the flagellins of Δ*maf-1* cells.

We conclude that Maf-1 and Maf-2 independently cause flagellins to migrate slower on SDS-PAGE, suggesting that both Mafs influence flagellin glycosylation in *S. oneidensis* MR- 1. By contrast, Maf-1 is a major determinant required for flagellar filament assembly, while Maf-2 contributes to filament length in the absence of Maf-1.

### Maf-1 and Maf-2 glycosylate flagellins on different serine residues

To directly show that Maf-1 and Maf-2 promote flagellin glycosylation and to determine which residues they modify (**Fig. 1E**), we conducted glycopeptide analyses by LC-MS/MS (liquid chromatography followed by tandem mass spectrometry)-based as established previously for FlaA/B of *S. oneidensis* MR-1^22^ (**Fig. 2A**). Since the polyclonal antibody to flagellins recognizes both FlaA/B, we individually expressed FlaA or FlaB harbouring an N-terminal 3xFLAG tag (FLAG-FlaA or FLAG-FlaB) from an IPTG-inducible *lac* promoter on pSRKGm^30^ in *WT* and *maf* mutant cells. Immunoblotting using monoclonal antibodies to the FLAG-tag revealed that loss of either Maf-1 and/or Maf-2 increases the mobility of FLAG-FlaA and FLAG-FlaB in *maf* single and double mutant cells compared to *WT* cells (**Fig. 2B**). These experiments essentially recapitulated the results seen in the immunoblots showed in **Fig. 1E** in which both flagellins are detected simultaneously using the polyclonal antibodies to FlaA/B.

**Figure 2.**
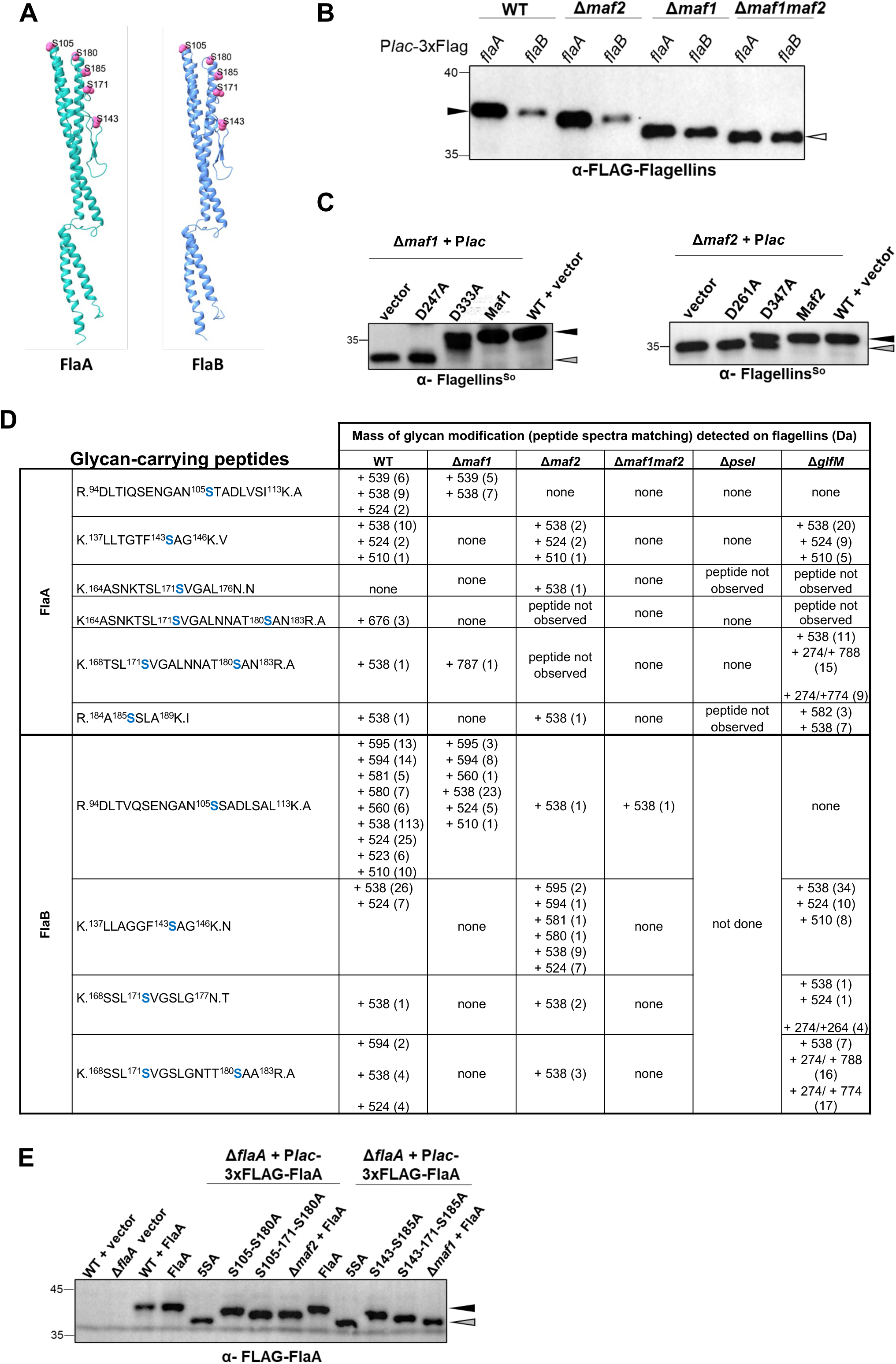
Maf-1 and Maf-2 are required for glycosylation of FlaA/B in *S. oneidensis* MR- 1. **A)** Structure prediction of FlaA/B by AlphaFold2 v2. The reported *O*-glycosylation serine sites (S105, S143, S171, S180 and S185) on both flagellins are indicated as pink spheres. **B)** Immunoblot analysis of 3xFLAG tagged FlaA and FlaB expressed from an IPTG- inducible P*_lac_* promoter on pSRK-Gm in *S. oneidensis WT* and mutant cells. Blots were probed with monoclonal antibodies to the FLAG-tag (α-FLAG Flagellins). The reduced migration position of glycosylated flagellin in *WT* cells is indicated by the black caret, while the white caret indicates the migration position of non-glycosylated flagellin in Δ*maf-1* Δ*maf-2* double mutant cells. **C)** Immunoblot analysis of FlaA and FlaB expressed in *S. oneidensis WT*, Δ*maf-1* (left) and Δ*maf-2* (right) single mutant cells expressing various versions of Maf-1 or Maf-2 from the IPTG-inducible P*_lac_* promoter on pSRK-Gm. Blots were probed with polyclonal antibodies to FlaA/B (α-Flagellins^So^). The black caret indicates the migration position of fully glycosylated, while the grey caret indicates the migration position of partially glycosylated flagellin in Δ*maf-1* and Δ*maf-2* single mutant cells. **D)** Glycopeptide LC-MS/MS analysis of flagellins FlaA and FlaB harbouring a 3xFLAG tag at the N-terminus. Flagellins were pulled down from *WT* and mutant cells extracts using anti-FLAG matrix and then subjected to glycopeptide analysis. Serine residues highlighted in blue correspond to the reported *O*-glycosylation sites of both flagellins ^22^ that are no longer detected in Pse-deficient (Δ*pseI*) cells. The listed peptides together with the excess masses were filtered based on the criteria of a score more than 200 and a mass error of 3 ppm. **E)** Immunoblot of 3xFLAG tagged FlaA and point mutant derivatives expressed from the IPTG-inducible P*lac* promoter on pSRK-Gm in *S. oneidensis WT* and Δ*flaA* cells. The black caret indicates the migration of *WT* FlaA complete glycosylation while the migration of mutants that are not or poorly glycosylated is indicated by the grey caret. Blots were probed with monoclonal antibodies to the FLAG-tag (α-FLAG-FlaA).

Previous glycopeptide analyses revealed five serine residues (S105, S143, S171, S180 and S185) bearing a 274 Da mass as base unit, likely corresponding to mono-*N*-acetyl- pseudaminic acid that is derivatized with an additional mass of either 236 Da, 250 Da (one methyl group on 236 Da moiety) or 264 Da (two methyl groups on 236 Da moiety), resulting in masses 510, 524 and 538 Da ^22^. Our LC-MS/MS-based glycopeptide analyses of FLAG- FlaA and FLAG-FlaB that we immunoprecipitated from *WT* cell extracts also revealed 510, 524 and 538 Da masses on the tryptic digests (**Fig. 2D**). Additionally, we detected masses above 538 Da in the FLAG-FlaA (i.e 676 and 787 Da) and FLAG-FlaB samples (i.e., 595, 594, 581, 580 and 560 Da) featuring the oxonium ion derived masses 275 Da and 257 Da of the 274 Da Pse version from *S. oneidensis* MR-1. Glycopeptide analyses of FLAG-FlaA and FLAG-FlaB extracted from Δ*maf1* and Δ*maf2* mutant cells (**Fig. 2D**) revealed a clear site-preference for Maf-1 *versus* Maf-2: in the absence of Maf-1, peptides carrying the expected glycosylation sites S105, S171 and S180 were detected for FLAG-FlaA, while only the peptide carrying the S105 modification were detected for FLAG-FlaB (**Fig. 2D**). By contrast, Maf-2 is not required for glycosylation of S143, S171 and S185 on FLAG-FlaA, while S105, S143, S171 and S180 are glycosylated on FLAG-FlaB (**Fig. 2D**). In contrast to previous reports^22^, we observed no modification on the S185-containing peptide of FLAG-FlaB extracted from *WT* cells (**Fig. 2D**).

These glycan modifications were barely detectable in the absence of Maf-1 and Maf- 2: no glycosylated peptides were detected on FLAG-FlaA and only one peptide (R^94^DLTVQSENGAN^105^SSADLSAL^113^KA) was detected carrying a modification with a 538 Da mass when FLAG-FlaB was extracted from Δ*maf-1*Δ*maf-2* cells. As this peptide was only detected once, it is possible that it a rare event stemming from promiscuous and inefficient glycosylation of FLAG-FlaB by a protein other than Maf-1 or Maf-2. It may also simply reflect a contamination from previous glycopeptide analyses or purifications. In summary, our glycopeptide analyses reveal a clear preference of Maf-1 for glycosylation on S143, whereas Maf-2 efficiently glycosylates S105.

A mutational strategy further corroborated this site-specificity of flagellin glycosylation detected by LC-MS/MS. To this end, we generated mutant derivatives of FLAG-FlaA with alanine substitutions at five serine residues simultaneously (S105, S143, S171, S180 and S185), at two serine residues simultaneously (S105-S180 or S143-S185) or at three serine residues simultaneously (S105-S171-S180 or S143-S171-S185). These versions were then expressed in Δ*flaA* cells and the glycosylation status of mutant FLAG-FlaA was subsequently determined by immunoblotting using antibodies to the FLAG tag. As seen in **Fig. 2E**, the alanine substitutions at three serine sites (S105-S171-S180) increase the mobility of FLAG- FlaA, thus mimicking the glycosylation status of FlaA in the Δ*maf-2* mutant. Unexpectedly, the combined alanine substitutions at S143-S171-S185 failed to mimic the glycosylation status of FlaA in the Δ*maf-1* mutant, suggesting that Maf-1 acts on (an) additional serine residue(s) of FlaA.

Since Maf-1 and Maf-2 both contain the MAF_flag10 domain, we predicted that they should both rely on a paralogous catalytic mechanism for glycosylation of their preferred serine acceptor sites. Indeed, mutation of the same aspartate residue to alanine (D247A and D261A for Maf-1 and Maf-2, respectively, **Supplementary Fig. S3A**), residue predicted to be required for function in Mafs ^20^, abrogated flagellin glycosylation in Maf-1 and Maf-2 and affected motility in the same manner as the gene disruptions (**Fig. 2C**, **Supplementary Fig. S3B, S3C**).

In summary, our results show that Maf-1 and Maf-2 are independently required for flagellin glycosylation in *S. oneidensis* MR-1, yet each with different preference for acceptor serine residues requiring a conserved aspartate residue in the MAF_flag10 domain.

### The C-terminal TPR domain of Maf-1 binds flagellin and is required for glycosylation

Having established that MAF_flag10 is required for activity, we determined whether the C-terminal TPR domain of Maf-1 is also required for flagellin glycosylation. To this end, we expressed a truncated Maf-1 version harbouring only the Maf_Flag-10 domain (1-420 aa) in Δ*maf-1* cells along with FLAG-FlaA or FLAG-FlaB from the same plasmid. Immunoblotting and motility assays revealed that truncated Maf1 (Maf1_1-420_) cannot complement the flagellin modification and motility defect of Δ*maf-1* cells. By contrast, when full-length Maf-1 is co-expressed with either FLAG-FlaA or FLAG-FlaB, these defects are reversed (**Fig. 3B-E**). We also constructed a truncated 1-699 version of Maf-1 lacking a C-terminal portion of the TPR and found that it only supports partial modification of both FLAG-FlaA (**Fig. 3B**) and FLAG- FlaB (**Fig. 3D**), but it does not confer motility to Δ*maf-1* cells. TEM analysis revealed that Maf- 1_1-699_ complemented cells expressing FlaA have shorter filaments as to an equivalent strain expressing FlaB in lieu of FlaA (**Fig. 3F**). Nonetheless, the filaments of these complemented cells are still shorter (**Fig. 3F**) compared to Δ*maf-1* cells complemented with full-length Maf-1 (**Fig. 1D**). This suggests that Maf-1_1-699_ is crippled in its ability to efficiently glycosylate flagellin.

**Figure 3.**
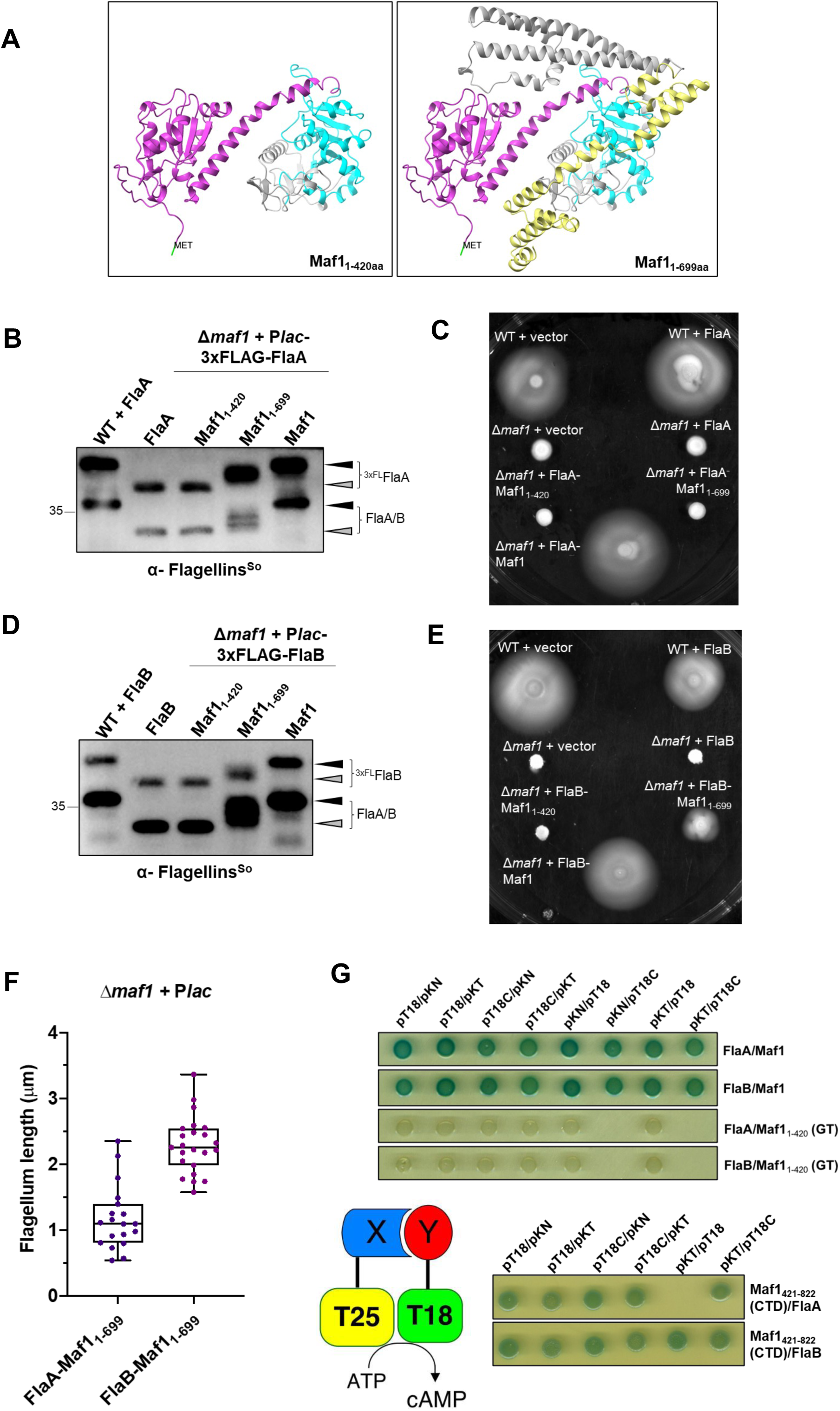
The CTD of Maf-1 is required for glycosylation and binding of FlaA/B. A) Structures of the truncated Maf-1_1-420_ and Maf-1_1-699_ versions predicted using ChimeraX. B) Immunoblot of 3xFLAG-FlaA and endogenous FlaA/B from *S. oneidensis WT*, Δ*maf*-*1* cells complemented with plasmids expressing various Maf-1 forms. FLAG-FlaA was co-expressed with Maf-1 or variants from P*_lac_* on pSRK-Gm or without Maf-1 just expressing FLAG-FlaA alone in *WT* and Δ*maf*-*1* cells (first two lanes from the left). Blots were probed using polyclonal antibodies to *S. oneidensis* FlaA/B (α-Flagellins^So^). C) Motility assay of *S. oneidensis WT*, Δ*maf*-*1*, and complemented strains on LB swarm (0.25%) agar. D) Immunoblot of 3xFLAG-FlaB and endogenous FlaA/B from *S. oneidensis WT*, Δ*maf*-*1* cells complemented with plasmids expressing various Maf-1 forms. FLAG-FlaB was co-expressed with Maf-1 or variants from P*_lac_* on pSRK-Gm or without Maf-1 just expressing FLAG-FlaB alone in *WT* and Δ*maf*-*1* cells (first two lanes from the left). Blots were probed with polyclonal antibodies to *S. oneidensis* FlaA/B (α-Flagellins^So^). E) Motility assay of *S. oneidensis WT*, Δ*maf*-*1*, and complemented strains on LB swarm (0.25%) agar. F) Flagellar filament length measurements in Δ*maf-1* cells harbouring a plasmid co- expressing Maf-1_1-699_ with FlaA and FlaB from P*_lac_*. G) Bacterial two hybrid assay in *E. coli* cells expressing T18 and T25 fusion proteins to *S. oneidensis* flagellins, full-length Maf-1, truncated Maf-1_1-420_ and the CTD of Maf-1_422- 822_. Cells were spotted on LB agar containing X-Gal (5-bromo-4-chloro-3-indolyl-β-D- galactopyranoside) and IPTG.

Next, we used bacterial two-hybrid assay (BACTH) to probe whether full-length Maf-1 or the TPR domain of Maf-1 binds FlaA/B directly. The BACTH assay which is based on the reconstitution of two fragments of adenylate cyclase (T18 and T25) from *Bordetella pertussis* in *E. coli* ^31^ grafted onto two proteins that interact directly. The readout for such *in vivo* reconstituted adenylate cyclase activity is LacZ induction in *E. coli.* Accordingly, we expressed full-length or truncated Maf1_1-420_ bearing T18 or T25 fusions from one plasmid and FlaA/B bearing the corresponding T18/T25 partner fragment from a compatible plasmid and then spotted the resulting clones on medium containing the LacZ substrate X-Gal (5-bromo-4- chloro-3-indolyl-β-d-galactopyranoside). As shown in **Fig. 3G**, all combinations of full-length Maf-1 with either FlaA or FlaB yielded LacZ-positive (blue) *E. coli* colonies. By contrast, when the TPR domain was deleted from Maf-1, colonies were no longer blue indicating that the deleted region is required for binding of FlaA/B. Indeed, we found that the CTD fragment of Maf-1 (Maf1_421-822_) that includes the TPR domain can support blue colony formation in this BACTH assay, indicating that the TPR alone can interact with flagellin (**Fig. 3H**).

We conclude that the TPR domain of Maf-1 binds flagellin and indispensable for flagellin glycosylation by Maf-1.

### GlfM promotes flagellin binding and glycosylation by Maf-2

Having established that the TPR domain identifies the correct acceptor for glycosylation by Maf-1, we next addressed the conundrum of how Maf-2 can specifically glycosylate its S105 target in the absence of an overt flagellin binding domain such as the TPR of Maf-1. Importantly, and consistent with the view that Maf-2 alone cannot bind flagellin, BACTH assays did not yield blue colonies when Maf-2 and FlaA/B fusions are co-expressed (**Fig. 4A**), suggesting that an unknown adaptor or specificity factor might act as bridge linking Maf-2 to FlaA/B. In examining the genetic environment of the Maf-2-encoding Gram-negative and Gram-positive bacteria whose flagellar *loci* encode both a putative Maf-2 orthologue and a Pse (or Leg) biosynthesis pathway, we noted a conserved coding sequence that flanks *maf- 2* (**Fig. 1A**, **Supplementary Fig. S2 and S4A**). In *S. oneidensis* MR-1, the corresponding *cis*- encoded gene product, SO_3260 (henceforth, <u>Gl</u>ycosylation factor for <u>M</u>af, GlfM, see **Fig. 1A**) is a soluble 89 residue protein predicted to assemble into a four-helix bundle according to AlphaFold modelling ^32^(**Fig. 4B**). Moreover, protein homology detection using the HHpred algorithm suggests that GlfM bears some sequence homology to the flagellar chaperones FliS or FliT. In support of the idea that GlfM acts as a Maf-2-specific bridging or adaptor protein, our analyses revealed that GlfM promotes the interaction between Maf-2 and FlaA/B in a tripartite BACTH system in which (untagged) GlfM is co-expressed in *E. coli* along with the tagged Maf-2 and FlaA/B fusion proteins bearing the T18 or T25 adenylate cyclase fragments (**Fig. 4A**). Since the classical bipartite BACTH analysis revealed that GlfM binds Maf-2, but neither FlaA/B nor Maf-1 (**Fig. 4A**), our BACTH experiments support the model that GlfM forms a ternary complex with Maf-2 and FlaA/B. They also suggest that by binding Maf-2, GlfM directs Maf-2 towards its acceptor (FlaA/B) to enable glycosyltransfer. To provide biochemical evidence for an interaction between GlfM and Maf-2, we conducted co-immunoprecipitation experiments of a FLAG-tagged GlfM variant (FLAG-GlfM) from extracts prepared from *S. oneidensis* MR-1 cells expressing FLAG-GlfM from pSRK-Gm. FLAG-GlfM and associated proteins were enriched by immunoaffinity chromatography using an anti-FLAG matrix and the pulled-down material was analysed by LC-MS/MS. We detected both FLAG-GlfM and Maf-2 in the pull-down samples (**Supplementary Fig. S4B**), indicating that GlfM and Maf-2 indeed form a complex in *S. oneidensis* MR-1.

**Figure 4.**
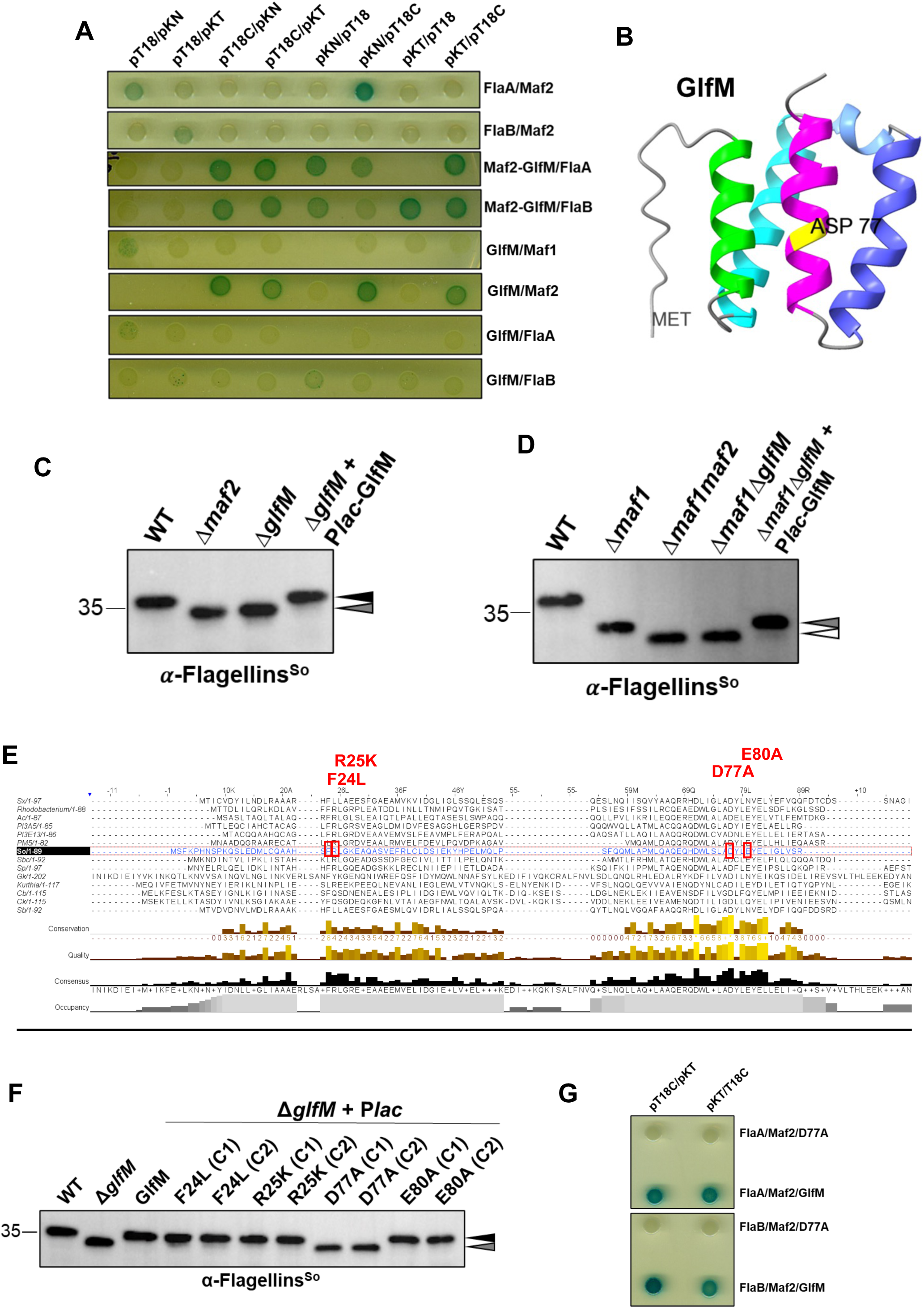
The GlfM accessory protein promotes flagellin binding and glycosylation. **A)** Bacterial two hybrid assay in *E. coli* cells expressing fusion proteins harbouring the T18 and T25 fragments from adenylate cyclase genetically encoded either at the N- or C-terminus of *S. oneidensis* flagellins, Maf-1 and Maf-2. GlfM was co-expressed from the same plasmid with the Maf-2 fusion protein. Cells were spotted on LB agar containing X-Gal and IPTG. **B)** Predicted structure of GlfM by AlphaFold v2 displaying four helices. The conserved and essential D77 is highlighted in yellow which lies at the last helix (in magenta). **C)** Immunoblot of *S. oneidensis* flagellins from *WT*, Δ*maf-2*, Δ*glfM* and complemented cells probed with antibodies to FlaA/B. The black caret indicates the migration position of glycosylated flagellin, while the grey caret denotes the position of partially glycosylated flagellin. Blots were probed with polyclonal antibodies to *S. oneidensis* FlaA/B (α-Flagellins^So^). **D)** Immunoblot of *S. oneidensis* flagellins in *WT*, Δ*maf-2*, Δ*maf-1Δmaf-2*, Δ*maf-1ΔglfM* and complemented cells. The grey caret indicates the migration position of partially glycosylated flagellin, while the white caret denotes the position of non-glycosylated flagellin. Blots were probed with polyclonal antibodies to *S. oneidensis* FlaA/B (α- Flagellins^So^). **E)** Multiple sequence alignment of GlfM orthologs using MUSCLE at EMBL-EBI. Sequence alignment is viewed using Jalview^56^. Point mutations (F24L, R25K, D77A and E80A) introduced in *S. oneidensis* MR-1 GlfM are indicated in red font. **F)** Immunoblot of *S. oneidensis* flagellins in extracts from *WT*, Δ*glfM* and complemented cells. Blots were probed with polyclonal antibodies to *S. oneidensis* FlaA/B (α- Flagellins^So^). The black caret indicates the migration position of glycosylated flagellin, while the grey caret denotes the position of partially glycosylated flagellin. **G)** Bacterial two hybrid assay of *E. coli* cells co-expressing *S. oneidensis* flagellins, and Maf-2 fused to T18 or T25. GlfM or point mutant GlfM-D77A was expressed separately from P*_lac_* on pSRK-Gm. Cells were spotted on LB agar containing X-Gal as described above for A.

If the interaction between GlfM and Maf-2 is required for the glycosylation of flagellin, then the loss of GlfM should phenocopy loss of Maf-2. To determine if this is the case, we generated a Δ*glfM* deletion mutant as well as a Δ*maf-1*Δ*glfM* double mutant in *S. oneidensis* MR-1 to compare the status of FlaA/B glycosylation to that of *WT*, Δ*maf-2* and Δ*maf-1*Δ*maf-2* cells by immunoblotting. The immunoblots shown in **Fig. 4C** indeed revealed that the migration of the flagellins in Δ*glfM* cells is identical to that of Δ*maf-2* cells and that this defect is corrected upon introducing a GlfM-expressing plasmid (pP*_lac_*-GlfM) into Δ*glfM* cells. Moreover, when comparing flagellin migration of Δ*maf-1*Δ*glfM* and Δ*maf-1*Δ*maf-2* double mutant cells (**Fig. 4D**) to that of the Δ*maf-1* parental cells, we noted that the Δ*glfM* mutation again has the same effect on flagellin glycosylation as the Δ*maf-2* mutation, at least by immunoblotting criteria. Again, this defect was reversed (to the Δ*maf-1* phenotype) when pP*_lac_*-GlfM was introduced into Δ*maf-1*Δ*glfM* cells. To confirm that GlfM is indeed required for Maf-2-dependent glycosylation of FlaA/B, we performed LC-MS/MS-based glycopeptide analyses of FLAG-FlaA and FLAG-FlaB extracted from Δ*glfM* cells. As seen in **Fig. 2C**, the modified peptides obtained for FLAG-FlaA and FLAG-FlaB closely correspond to the mass detected when flagellin that had been extracted from Δ*maf-2* cells was analysed.

Since GlfM orthologs are found both in Gram-negative and Gram-positive bacteria, we aligned the GlfM sequences to identify conserved residues that could be required for GlfM function. A multiple sequence alignment in GlfM orthologs (**Fig. 4E**), revealed F24, R25, D77 and E80 as strictly conserved residues. We then conducted site-directed mutagenesis of these residues to test whether GlfM variants with single substitutions can still complement the glycosylation defect of *S. oneidensis* MR-1 Δ*glfM* cells when expressed from pP*_lac_*-GlfM. Immunoblotting (**Fig. 4F, Supplementary** Fig. 4C) revealed that the D77A mutant no longer supports GlfM function, whereas the substitution of F24, R25 and E80 did not cripple GlfM activity. D77 is predicted to reside in the last α-helix of GlfM (**Fig. 4B**), suggesting that the last helix promotes the interaction between GlfM and Maf-2. Indeed, BACTH assays revealed that GlfM-D77A no longer enables the Maf-2 interaction with FlaA or FlaB (**Fig. 4G**).

Taken together, our analyses show that unlike Maf-1, Maf-2 is dependent on the *cis*- encoded and conserved GlfM protein to bind and glycosylate flagellins. Moreover, D77 and/or helix 4 perform a key role in the function of GlfM and likely its orthologs.

### GlfM and Maf-2 constitute a tripartite glycosylation system

Since glycosylation by Maf-2 in *S. oneidensis* MR-1 requires at least three protein components (Maf-2, GlfM and FlaA/B), we revisited our earlier result that co-expression of Maf-2 and flagellin did not alter flagellin migration in *C. crescentus WT versus* Δ*neuB* cells, especially after having independently confirmed that flagellin glycosylation in *S. oneidensis* MR-1 depends on the Pse synthase PseI (**Fig. 5A**). We therefore designed a three-component co-expression system including Maf-2, flagellin and GlfM in *C. crescentus WT* and Δ*neuB* cells. We expressed either pP*_lac_*-*flag-flaA-maf-2* or pP*_lac_*-*flag-flaB-maf-2* for co-expression of FLAG-FlaA or FLAG-FlaB and Maf-2 from P*_lac_*. In a subsequent step, we introduced a second, compatible plasmid (pP*_xyl_*-*glfM*) expressing GlfM from the xylose-inducible P*_xyl_* promoter or the empty vector (pMT375) into these cells and then probed for decreased mobility of FLAG-FlaA or FLAG-FlaB by immunoblotting using antibodies to the FLAG tag. As shown in **Fig. 5B**, we observed a decrease of mobility of FLAG-FlaA/B when cells contain Pse, GlfM and Maf-2. In the absence of the GlfM-expression plasmid, the mobility of FLAG-FlaA or FLAG-FlaB is clearly increased.

**Figure 5.**
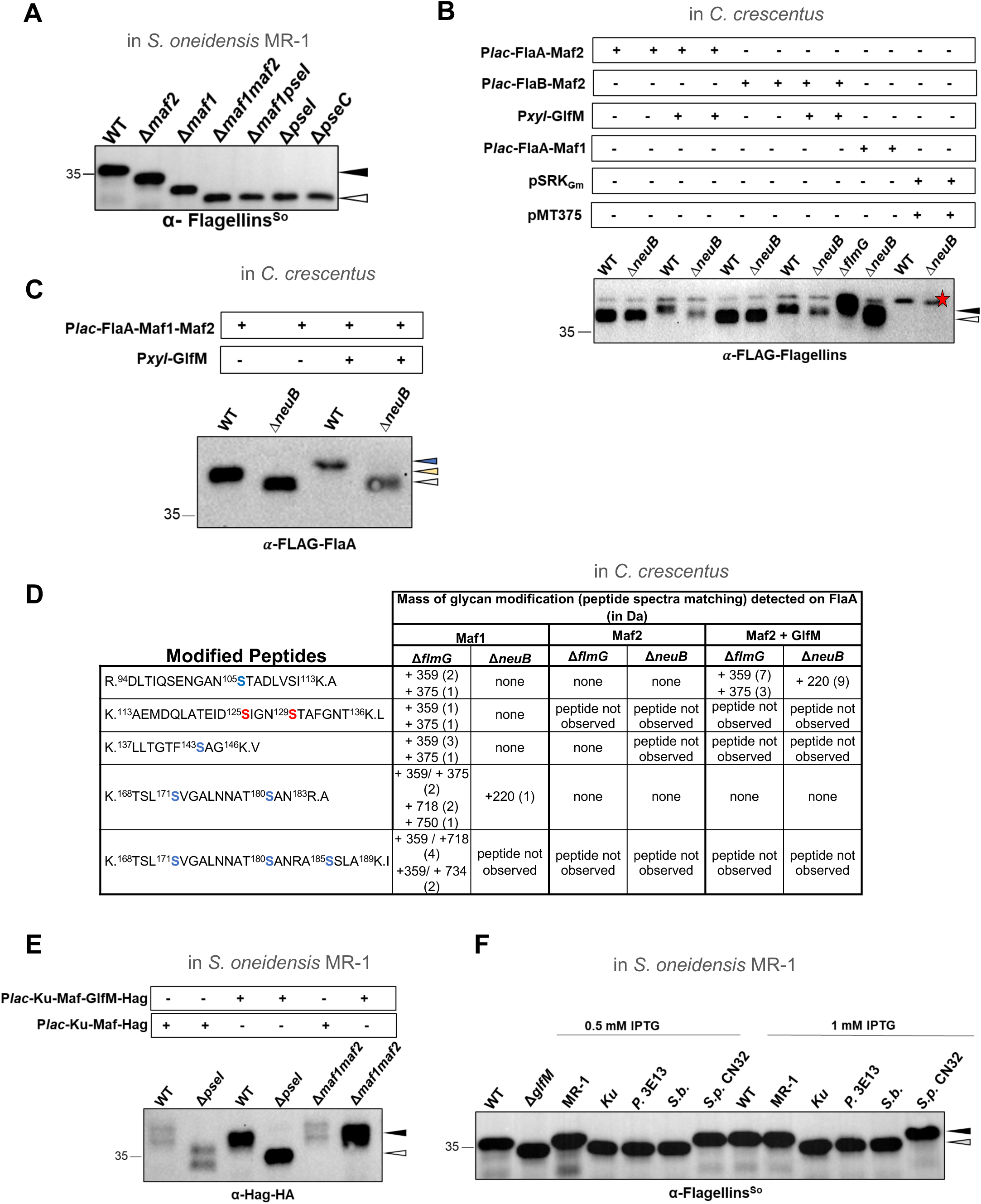
Reconstitution of different Maf-2 and GlfM pairs in heterologous hosts. **A)** Immunoblot of *S. oneidensis* flagellins in *WT*, Δ*maf-1*, Δ*maf-2*, Δ*maf-1*Δ*maf-2*, Δ*maf- 1*Δ*pseI*, Δ*pseI* and Δ*pseC* cells. Black caret indicates complete glycosylation while white caret indicates complete loss of modification. Blots were probed with polyclonal antibodies to *S. oneidensis* FlaA/B (α-Flagellins^So^). **B)** Immunoblot of *S. oneidensis* FLAG-tagged flagellins expressed in *C. crescentus WT*, Δ*flmG* and Δ*neuB* in the presence or absence of MR-1 GlfM. Black caret indicates complete glycosylation while, the white caret indicates complete loss of modification. Blots were probed with monoclonal antibodies to the FLAG-tag (α-FLAG-Flagellins). The red star denotes a cross-reacting protein. **C)** Immunoblot of *S. oneidensis* FLAG-tagged FlaA expressed in *C. crescentus WT* and Δ*neuB* in the presence or absence of *S. oneidensis* MR-1 GlfM. The yellow caret indicates the reduced migration of FLAG-FlaA, while the blue caret indicates the intermediate migration of FLAG-FlaA and the white caret indicates the complete loss of modification. Blots were probed with monoclonal antibodies to the FLAG-tag (α- FLAG-FlaA). **D)** Glycopeptide LC/MS-MS analysis of 3xFLAG-tagged FlaA purified from *C. crescentus* Δ*flmG* and Δ*neuB* co-expressing either FLAG-FlaA and Maf-1, FLAG-FlaA and Maf-2 or FLAG-FlaA, Maf-2 and GlfM. Serine residues that are in blue correspond to the reported *O*-glycosylation sites of FlaA/B^22^. None indicates the absence of glycan modification on the peptides that are glycosylated in *WT* cells. The listed peptides together with the excess masses were filtered based on the criteria of a score more than 200 and a mass error of 3 ppm. **E)** Immunoblot of HA-tagged Hag flagellin (Hag-HA) of *Kurthia* sp. 11kr321 expressed in *S. oneidensis* WT, Δ*pseI* and Δ*maf-1Δmaf-2* in the presence or absence of the cognate GlfM of *Kurthia*. The black caret denotes the migration of completely glycosylated Hag- HA, while white caret indicates the migration of non-glycosylated Hag-HA. Blots were probed with monoclonal antibodies to the HA-tag (α-Hag-HA). **F)** Immunoblot of flagellins in *S. oneidesis WT* and Δ*glfM* cells complemented cells, either expressing MR-1 GlfM (MR-1) or GlfM orthologs from *Kurthia* sp. 11kr321 (*Ku*), *Pseudomonas* sp. Irchel 3E13 (*P. 3E13*), *Shewanella baltica* (*S. b.*) and *Shewanella putrefaciens* CN-32 (*S. p. CN32*). Blots were probed with polyclonal antibodies to *S. oneidensis* FlaA/B (α-Flagellins^So^).

Yet, we also noted an increase in mobility when cells are unable to produce the Pse glycosylation donor, albeit here the change in migration is less pronounced compared to that seen in the absence of GlfM. As additional control, we also transformed single co-expression plasmid (pP*_lac_*-*flag-flaA-maf-1*) encoding FLAG-FlaA and Maf-1 into *C. crescentus* Δ*flmG* cells (lacking the native FlmG-type fGT) and Δ*neuB* cells as a reference point ^16^ to show the migration of flagellin after Pse-dependent glycosylation by Maf-1 in *C. crescentus* (**Fig. 5B**). Since the flagellins are glycosylated by both Maf-1 and Maf-2 in *S. oneidensis* MR-1, we also reconstituted both Maf activities in *C. crescentus* by first transforming *WT* and Δ*neuB* cells with a tri-cistronic co-expression plasmid encoding FLAG-FlaA, Maf-1 and Maf-2 (pP*_lac_*-*flag- flaA-maf-1-maf-2*) and then introducing the pP*_xyl_*-*glfM* supporter plasmid for Maf-2 activity. Immunoblotting using antibodies to the FLAG tag revealed that the presence of both Mafs further decreased the mobility of FLAG-FlaA compared to *C. crescentus* cells that are only featuring Maf-1 activity (**Fig. 5C**), a result that mirrors the joint contribution of Maf-1- and Maf- 2-dependent glycosylation to flagellin mobility in *S. oneidensis* MR-1 (**Fig. 4D**).

Next, we extracted FLAG-FlaA from the reconstituted *C. crescentus* strains using the anti-FLAG affinity matrix and subjected the samples to glycopeptide analysis (**Fig. 5D**). LC-MS/MS analyses revealed that S105 is indeed glycosylated by Maf-2, but only when GlfM is present. We did not detect any Maf-2-dependent glycosylation on any residues on FLAG-FlaA in *C. crescentus* lacking GlfM. The Maf-2-GlfM-dependent mass modifications are 359 Da or 375 Da, corresponding to the di-acetylated amide and oxidized forms of Pse specific to *C. crescentus* (also detected in our previous analyses of FlmG-dependent glycosylation)^16^. By contrast, when FLAG-FlaA was extracted from Δ*neuB C. crescentus* cells that do not produce Pse (derivatives), the 359 Da and 375 Da mass were no longer detected. Instead, LC-MS/MS revealed an alternative added mass of 220 Da on FLAG-FlaA that likely stems from the addition of a KDO moiety (3-deoxy-manno-octulosonic acid), an 8-carbon acidic sugar from the inner core of lipopolysaccharide that can be mistakenly donated by Mafs when the preferred nonulosonic acid (Pse or Leg) is not available^33^. When we analysed glycosylation of FLAG-FlaA by Maf-1 in *C. crescentus*, the 359 Da and 375 Da modifications were found on the peptides with S143, as well as those with S171, S181, S185, S125, S129 or S105.

Our results confirm that Maf-2 and GlfM are both required for site-specific glycosylation of flagellin with Pse, whereas Maf-1 can glycosylate flagellin autonomously. Whereas Maf-1 requires its C-terminal TPR domain for glycosylation, it appears that its activity is more promiscuous on the acceptor residue, at least in the heterologous host *C. crescentus*, compared to the glycosylation reaction executed by the GlfM-Maf-2 system.

### Conservation of the GlfM-Maf-2 glycosylation module in Gram-negative and Gram-positive bacteria

Genome comparisons of flagellar gene clusters containing Pse/Leg biosynthesis revealed GlfM-like coding sequences next to the genes encoding Maf-2 orthologs (**Fig. 4E**, **Supplementary Fig. S2 and S4A**). This genetic architecture suggests that Maf-2 orthologs depend on GlfM for function. To investigate whether GlfM also serve a similar function in Gram-positive lineages, we selected the GlfM-Maf2-Hag (note that the flagellin is called Hag) system of the Gram-positive bacterium *Kurthia sp.* 11kri321 for two important reasons. First, its flagellin is glycosylated at several serine and threonine residues^15^. Second, its Maf-2 and GlfM are encoded just adjacent to the Pse biosynthesis genes^16^ within the flagellar gene cluster. Co-expression plasmid either with GlfM (pP*lac*-*maf*-*glfM*-*hag*) or without (pP*lac*-*maf*- *hag*) were mobilized into *S. oneidensis WT*, Δ*pseI* and Δ*maf-1*Δ*maf-2* double mutant cells. Immunoblotting of an HA-tagged *Kurthia* Hag (Hag-HA) co-expressed with *Kurthia* GlfM and Maf-2 in *S. oneidensis* MR-1 Δ*maf-1*Δ*maf-2* double mutant cells revealed glycosylation of Hag-HA as judged by its change in mobility (**Fig. 5E**) that is Pse-dependent since Hag-HA appeared non-glycosylated in Δ*pseI* cells that lack Pse. Importantly, accumulation of this form of Hag-HA was strongly dependent on the presence of GlfM (**Fig. 5E**). Similarly, expression of *Kurthia* Maf-GlfM and its cognate flagellin in *C. crescentus* showed that GlfM promotes the accumulation of glycosylated Hag-HA (**Supplementary Fig. S4D**). Finally, we confirmed that the Maf of *Pseudomonas* sp. 3E13 also requires GlfM to support glycosylation of its cognate flagellin in *C. crescentus* (**Supplementary** Fig. 4E).

Since cognate GlfM-Maf-2-flagellin pairs from Gram-positive and Gram-negative lineages function together, we determined the specificity of GlfMs by testing whether they can promote glycosylation by noncognate Maf-2. To this end, we conducted complementation of *S. oneidensis* MR-1 Δ*glfM* cells with selected GlfM orthologs from Gram-negative and Gram-positive bacteria and found that only the GlfM ortholog from *Shewanella putrefaciens* CN-32 can restore flagellin mobility to *S. oneidensis* MR-1 Δ*glfM* cells (**Fig. 5F**). This *S. putrefaciens* CN-32 GlfM ortholog can also functionally interact with Maf-2 from *S. oneidensis* MR-1 to promote flagellin glycosylation in the heterologous host *C. crescentus* (**Supplementary Fig. S4F**). Interestingly, the GlfM ortholog from *Shewanella baltica* did not functionally interact with Maf-2 from *S. oneidensis* MR-1 in both assays (**Fig. 5F, Supplementary Fig. S4F**). Knowing that the *S. baltica* Maf-2 is linked to a putative Leg biosynthesis cluster, we speculate that the GlfM-Maf-2 systems may also exhibit glycosyl donor specificity akin to the FlmG orthologs.

## DISCUSSION

Despite the prevalence of glycosylated flagella in the bacterial domain of life, unambiguous proof showing that the conserved Mafs are indeed flagellin sialyltransferases in Gram-negative and Gram-positive bacteria had not been provided. By demonstrating sufficiency for two paralogous, yet mechanistically distinct, Maf systems from *S. oneidensis* MR-1, we prove that these widely conserved enzymes are protein sialyltransferases, in this case executing flagellin O-glycosylation with a Pse derivative as donor. Remarkably, we discovered that different Mafs can exhibit exquisite target serine preference on the same acceptor. Consistent with the notion that different modes of acceptor binding define this target serine selection, we discovered that Maf-1 and Maf-2 use different types of flagellin binding domains. While the C-terminal TPR domain in Maf-1 binds flagellin and resides within the same polypeptide, we uncovered a conserved *cis*-encoded four-helix bundle protein, GlfM, that promotes ternary complex formation between Maf-2 and flagellin. In both cases, these domains are critical for Maf-dependent glycosylation, indicating that complex formation between Maf and flagellin is required for glycosylation activity. We propose that such distinct binding modes to flagellin could place different glycosyl accepting serine in the catalytic centre, ultimately dictating which serine is best positioned as acceptor OH group for each Maf.

It is also possible that a different active site intrinsic to the GT domain architecture of Maf-1 *versus* Maf-2 determines which serine residue in the flagellin is accessible as acceptor for Pse, independently of which flagellin binding domain is used. Mafs differ from the FlmG- class of fGTs in the presence of a GT-A type GT domain, whereas the FlmGs harbour a GT- B type domain^5^. The GT-A fold typically relies on two nearby aspartate residues for coordination of a metal ion that is critical for catalysis^17,34^. Our work provided evidence that the orthologous aspartate residue is required for function in Maf-1 and Maf-2, akin to the corresponding Maf point mutation in *M. magneticum* AMB-1^20^. Structural determination with flagellin will clarify how the accepting residue is placed in the GT domain active site and whether the organisation of the catalytic centre differs substantially between Maf-1 and Maf-2. It will also be interesting to compare at the structural level how flagellin is bound by Mafs *versus* FlmG, and contrast this mode of binding to that of other flagellin binding proteins, such as flagellin secretion chaperones or translational regulators of flagellin mRNAs ^35–39^. While these flagellin binding proteins reside in the cytoplasm, there are others such as the innate immune receptor TLR5 ^40,41^ or flagellin capping proteins that must bind flagellin in the extra- cytoplasmic space^11^. Capping proteins or analogous flagellin polymerization factors^12^ act as chaperones that promote the insertion of flagellin at the distal end of the flagellar filament, once the unfolded flagellins emerge at the end of the secretion tunnel inside the flagellar filament following their export from the bacterial cytoplasm ^11,12,42^. To ensure that they are located at the distal end of the nascent filament before the flagellins arrive, capping proteins are thought to be secreted through the flagellar translocation apparatus before the flagellins (are expressed)^12^. By contrast, flagellin can only bind TLR5 in the extracellular space upon dissociation from the filament or upon escape from the capping/polymerization protein. With several flagellin binding proteins residing in the cytoplasm, the question of whether and how these interactions might occur sequentially to prevent competitive binding by multiple proteins at the same time will need to be investigated in future work. An intriguing possibility posits that their relative affinities for flagellin differ before or after glycosylation has occurred, albeit it is important to recall that not all flagellated bacteria modify flagellin with Pse or Leg.

### Conserved function and evolution of GlfM-Maf-2 systems

In FlmG, the C-terminal domain dictates whether Pse or Leg is transferred, while the N-terminal TPR domain binds flagellin. Maf-1 uses a C-terminally located TPR to execute the analogous function, but Maf-2 lacks such a C- or an N-terminal extension and it cannot bind flagellin on its own. The *cis*-encoded and conserved GlfM proteins has been appropriated to serve as bridging protein for Maf-2 to bind flagellin and facilitate glycosylation. Therefore, while Maf-1 and FlmG are fundamentally two-component protein sialyltransferase systems, the GlfM-Maf-2 class is the first example of a tripartite system. We suspect that GlfM could actively contribute to contact points for direct interactions with flagellin while simultaneously binding Maf-2 using other residues. It is also possible that GlfM induces a structural change in Maf-2 that allows residues from Maf-2 to wrap around flagellin, such that GlfM does not physically associate with flagellin itself.

We used several heterologous hosts to demonstrate that GlfM and Maf from Gram-negative and Gram-positive bacteria act together. Sequence alignments with GlfM orthologs and subsequent genetic analyses using GlfM from *S. oneidensis* MR-1 as model revealed the conserved D77 residue as a key determinant for GlfM in promoting an interaction between Maf and flagellin. It will be interesting to determine whether the aspartate corresponding to D77 in other GlfM orthologs Gram-positive bacteria is also required for Maf-2 activity. It may not be coincidental that GlfM generally seems to be encoded downstream of the Maf-2 coding sequence in Gram-negative and Gram-positive genomes. In fact, it is tempting to view this bi- cistronic organization as a relic of ancestral Maf-1-type composed of a single polypeptide that once included a C-terminal flagellin binding domain by a GlfM domain that eventually became separated during evolution by the acquisition of stop/start codons that gave rise to individual translation products. In an alternative evolutionary path, originally separate Maf-2 and GlfM coding sequences could have become joined to encode within a single Maf-1-type fusion protein. A single polypeptide in which a Maf domain is joined to a C-terminal flagellin binding domain would ensure efficient glycosylation once flagellin is bound and would eliminate possible stoichiometric fluctuations between flagellin-binding and glycosyltransferase domains. By contrast, a two-protein combo may offer regulatory versatility, for example with GlfM potentially serving multiple Mafs or regulating other proteins that depend on direct interactions with flagellin. Indeed, there are bacterial genomes such as *Campylobacter jejuni* that encode numerous Maf orthologs and that produces three different sialic acid derivatives ^1,3,18^. It is also conceivable that GlfM could be used to control proteins that do not affect motility. In this case, GlfM could serve to coordinate flagellin glycosylation with other cellular events.

It has not escaped our attention that GlfM and the TPR of Maf-1 are both predicted to form four-helix bundles, raising the possibility that binding of flagellin by fGTs occurs through similar alpha-helical bundles. From the functional and structural view point, GlfM shares features with two-component signalling (TCS) systems that also bear four-helix bundles ^43^, or possibly the TCS connector proteins ^44,45^. TCS connectors are small ancillary proteins that tune the activity of histidine kinases (HKs) and their ability to phosphorylate their cognate response regulators (RRs). Akin to the direct and highly specific HK-RR interaction needed for phosphorylation of the RR acceptor, fGTs must also interact directly and exclusively with their cognate flagellin for the post-translational modification of the acceptor site. Moreover, since a four-helix bundle also underlies the operational management of HK-RR systems, the additional complexity of fGT-flagellin systems lies in the variation of the donor glycan that is transferred by fGTs as compared to the standard ATP from which the donating phosphate group stems in HK-RR systems^46–48^.

### Sialic acid donor specificity of Maf-1 and Maf-2

In the *S. oneidensis* MR-1 genome, the Maf-1 and Maf-2 coding sequences flank the coding sequences for Pse biosynthesis proteins and several other proteins of unknown function that may act to produce a substantially modified version of Pse^22,27^. We have shown that Maf-1 and Maf-2 can glycosylate flagellin with the *C. crescentus* Pse, a derivative having a mass of 359 or 375 Da that is used as glycan modification for flagellin, rather than the 317 Da mass that is predicted for the standard Pse. Nonetheless, this Pse mass is also smaller than the modification typically detected on flagellin of *S. oneidensis* MR-1, most frequently a mass of 538/9 or 524 Da. Therefore, Maf-1 and Maf-2 probably possess a preference for the large-mass Pse derivative of *S. oneidensis* MR-1.

Having recently demonstrated for the FlmG class that Pse-specific and Leg-specific fGTs exist in *C. crescentus* and in *B. subvibroides*^4,16^, respectively, we speculate that Maf proteins preferring Leg over Pse as substrate also exist for the Maf class. In fact, while the orthologous *locus* of *S. putrefaciens* CN-32 encodes Pse biosynthesis proteins, others such as *Shewanella sp.* KJ2020 (Genbank: JAKUKD010000032.1), *Shewanella baltica* OS185 (GenBank: CP000753.1) and *Shewanella xiamenensis* DSM22215 (GenBank: JAKILI010000004.1) likely encode Leg biosynthesis proteins flanked by the GlfM-Maf-2 and Maf-1 coding sequences. A hallmark of the Leg-biosynthesis pathway is the presence of the LegX enzyme which is required for the conversion of the UDP-linked N-acetylglucosamine (UDP-GlcNAc) to GDP-GlcNAc, the starting point for Leg biosynthesis^16^. In fact, L2745_04770, the *S. xiamenensis* DSM22215 homolog of *B. suvibrioides* LegX (Bresu_3267) is encoded upstream of the corresponding Leg-cytidylyltransferase and the Maf-1 ortholog. Pse and Leg, but also KDO, are activated by specialized cytidyltransferases that add a CMP- group to the precursor to make it a suitable substrate for the fGTs in the case of a protein acceptor or other GTs in the case of glycan-acceptor in O-antigen or capsule synthesis. While the corresponding cytidyltransferase encoded in the *S. oneidensis* MR-1 Pse biosynthetic operon (SO_3269, **Fig. 1A**) has not yet been genetically analysed, it is much larger than the canonical cytidyltransferases, perhaps pointing towards a specialized reaction that it executes to produce the atypical Pse-derivative of *S. oneidensis* MR-1 with mass 538/9 or 524.

The Maf-2 (GK_3129) and GlfM (GK_3130) orthologs from the Gram-positive bacterium *Geobacillus kaustophilus* are also embedded in a flagellar locus containing a nearby LegX (GK_3121) coding sequence^33^. When heterologous expression of the Maf-2 ortholog was attempted in a sialic acid producing *E. coli*, the GlfM ortholog (GK_3130) had not been co-expressed with flagellin and Maf-2. Only a mass corresponding to KDO was rarely added to flagellin ^33^, but efficient transfer of sialic acid did not occur. We speculate that this failure likely can be attributed to two omissions. Firstly, the absence of the critical specificity factor GlfM and, secondly, the lack of a suitable Leg-derived donor sialic acid for this Maf-2 derivative.

In conclusion, having established sufficiency with bipartite and tripartite Maf systems, the current goal will be to attempt re-wiring the sialic acid donor of Mafs to broaden the repertoire of cytoplasmic protein glycosylation systems that can be used for future biomedical or biotechnological applications^49–51^.

## ACKNOWLEDGMENTS

We are grateful to Kai Thormann (University of Giessen, DE) for providing materials, and to Kai Thormann and Gert Bange (University of Marburg, DE) for discussions on Maf-1. We are very indebted to Dr. Chia-wei Lin and Dr. Sybille Pfammatter from the Functional Proteomics Center of Zürich for outstanding technical support with the flagellin glycopeptide and protein identification analyses, respectively. This work was funded by Swiss National Science Foundation (SNSF) project grant 310030_212531 (to P.H.V).

## AUTHOR CONTRIBUTIONS

Conceptualization: J.U., N.K. and P.V.; Methodology: J.U., N.K. and P.V; Validation: J.U., N.K. and P.V.; Formal analysis: J.U., N.K. and P.V; Investigation: J.U., and N.K.; Resources: J.U., N.K. and P.V.; Writing – Original Draft: J.U. and P.V.; Writing – Review & Editing: J.U., N.K. and P.V.: Visualization: J.U., N.K. and P.V.; Supervision: P.V.; Funding Acquisition: P.V.

## EXPERIMENTAL PROCEDURES

### Bacterial strains and culture conditions

Strains used in this study are listed in **Supplementary Table S1**. *Shewanella oneidensis* MR-1 and *Escherichia coli* recombinants were grown in/on Lysogen Broth (LB) medium at 30°C and 37°C, respectively. *Caulobacter crescentus* was grown in peptone-yeast extract (PYE) medium at 30°C as previously described (Kint et al 2023). When needed, medium (liquid/solid) was supplemented with antibiotics at the following final concentrations: for *S. oneidensis* transconjugants, 5 μg/mL gentamicin and 25 μg/mL of and kanamycin, respectively; for *C. crescentus* transformants, 1 μg/mL of gentamicin and 1 μg/mL of tetracyline; for *E. coli* transformants, 10 μg/mL of gentamicin and 10 μg/mL of tetracycline, 25 μg/mL or 50 μg/mL of kanamycin, and 120 μg/mL or 200 μg/mL of ampicillin. For conjugation using the auxotroph *E. coli* WM3064 donor, LB medium was supplemented with 0.3 mM meso- 2,6-diamino-pimelic acid (mDAP). LB plates containing 10% sucrose (w/v) were used for SacB-mediated counterselection of double recombinants. Isopropyl-beta-D-thiogalactosidase (IPTG) was used at 0.5 mM to induce gene expression from the *lac* promoter (P*_lac_*) on pSRKGm-derived plasmids in *C. crescentus* and *S. oneidensis*, while at 1 mM used for induction of P*_lac_* in *E. coli*. Additionally, xylose was used at 0.03-0.06% for induction of gene expression from the xylose-inducible (P*_xyl_*) promoter in *C. crescentus*.

Conjugations for intergeneric plasmid transfer from *E. coli* to recipient cells were set- up by individually washing *E. coli* WM3064 harbouring the donor plasmid and recipient *S. oneidensis* cells in LB, then mixing them at 1:10 ratio, followed by incubation for 2-4 hours at 30°C on LB plates harbouring mDAP.

For motility assays, 1 μL of *S. oneidensis* cultures previously grown for 4 hours to OD_600_=1.0 were spotted on semi-solid LB plates containing 0.25% agar and the appropriate antibiotics and IPTG. Motility was scored after 2 days incubation at 30°C by photography.

### Immunoblotting

*S. oneidensis* or *C. crescentus* were grown for 4 hours to OD_600_=1.0 or 0.4, respectively. Cells were harvested by centrifugation and cell pellets were used for immunoblot analyses. Lysates were prepared by first resuspending cell pellets in Laemmli buffer (Laemmli, 1970), followed by boiling at 95°C for 10 minutes. Cell-free extracts were then resolved by SDS-PAGE using 12.5%-15% polyacrylamide gels and subsequently electro-blotted onto a 0.45 μm polyvinylidene difluoride (PVDF) Immobilon®-P membrane (Merck Millipore, Sigma Aldrich) at 100 V for 90 minutes. The blot was then blocked with 5% milk in 1X TBS buffer supplemented with 0.1% Tween 20. For the detection of Flag-tagged flagellins, OctA-Probe monoclonal antibody (at dilution 1:1000; Santa Cruz Biotechnology) or anti DYKDDDK tag monoclonal antibody (at dilution 1:1000; Proteintech) were used. For detection of untagged *S. oneidensis* flagellins, polyclonal antibodies (at dilution 1:4000) against FlaA/B (Bubendorfer et al. 2013) were used. For the detection of HA-tagged flagellin, a monoclonal HA antibody (at dilution 1:4000, Proteintech) was used. Horseradish peroxidase-conjugated secondary antibodies (anti-mouse or anti-rabbit, at dilution 1:10000) and Immobilon® Western Chemiluminescent HRP substrate (Merk Millipore, Sigma Aldrich) were used to visualize FLAG-FlaA/FlaB and untagged FlaA/FlaB.

### Transmission electron microscopy and shearing of flagella

For transmission electron microscopy (TEM), *S. oneidensis* cells were negatively stained with uranyl acetate using a previously described protocol^12,16^. Briefly, cells were harvested after 4 hours of growth and resuspended in sterile water. Prior to depositing 20 μL of resuspended cells onto the 200-mesh copper, carbon-coated formvar grids (EM Science, Hatfield, PA), the grids were glow discharged for 1 minute. Grids were then allowed to adsorb the cells for 1 minute, followed by three consecutive washes in water, staining with 1% uranyl acetate for 1 minute and lastly, washing with water for 30 seconds. Negatively stained cells on grids were then imaged using a Tecnai 20 (FEI Company, Eindhoven, Netherland) electron microscope. Flagellum length measurements were carried out using the ImageJ software^52^.

For shearing *S. oneidensis* flagellum, 10 mL of cultures grown for 4 hours were harvested and resuspended in 200 μL LB medium, followed by gently passing cells through a syringe-needle for 5 minutes. After centrifugation at 12000 rpm for 1 minute, supernatant was then collected as the sheared fraction.

### Co-immunoprecipitation, glycopeptide analysis and protein identification

Flagellins were immunoprecipitated from extracts prepared from *S. oneidensis* or *C. crescentus* cells harvested after 4 of growth. While cell lysis of *C. crescentus* was done using Ready-Lyse™ and Nonidet P (NP)-40, as described in^16^, *S. oneidensis* cells were lysed only by boiling at 95°C for 15 min. Extracts were then treated with The DYDDDDK Fab-Trap™ agarose beads (ChromoTek, Proteintech) and the retained Flag-tagged FlaA/B was eluted from the agarose beads using 3xFLAG®-peptide (Chromotek, Proteintech). For the co-IP of GlfM from *S. onedensis* cell extracts, cell lysis was done using Ready-Lyse™, Lysozyme and NP-40. GlfM was also immunoprecipitated using DYDDDDK Fab-Trap™ agarose beads (ChromoTek, Proteintech) and eluted using 2X Laemmli buffer. Eluted flagellins and GlfM were then sent to Functional Genomics Center at the University of Zurich for glycopeptide analysis and protein identification by liquid chromatography-tandem mass spectrometry (LC-MS/MS), respectively.

For glycopeptide analyses, 10 μg of samples were digested with 500 ng sequence-grade trypsin (Promega) overnight at 37°C. Subsequently, the digested samples were dried, dissolved in 20 μL double-distilled water with 0.1% formic acid and transferred to the autosampler vials for LC-MS/MS. Two microliters of samples were injected on a nanoAcquity UPLC coupled to an Orbitrap Fusion™ mass spectrometer (ThermoScientific). Data analysis was done using the Byonic™ software with the following search parameters: mass tolerance of 10 ppm and fragment ion tolerance of 0.03 Da. Glycan wildcard search was done by considering mass adducts between 130-600 or 130-900 Da on serine residues.

For protein identification, gel bands were washed twice with a mixture of 100 mM NH_4_HCO_3_/50% acetonitrile followed by a wash with acetonitrile. Gel pieces were then digested with trypsin in buffered solution at pH 8.0 (10 mM Tris, 2mM CaCl_2_). Digested samples were collected, dried, and dissolved in aqueous solution of 3% acetonitrile with 0.1% formic acid. Peptides were separated on a M-class UPLC and analysed on a Orbitrap Fusion™ mass spectrometer (Thermofischer). MS data were processed using PEAKS Studio Plus (Bioinformatic Solutions) and spectra were searched against *Shewanella oneidensis* MR-1 protein database. Results were viewed using the Scaffold 5 viewer (Proteome Software).

### Bacterial adenylate cyclase two-hybrid (BACTH) assays

The entire open reading frames (ORsF) of *flaA*, *flaB*, *glfM*, *maf1*, *maf2*, as well as the N-terminal and central domain (GT, amino acids 1-420) and C-terminal (TPR, amino acids 421-822) parts of *maf1* were fused either with T18 or T25 fragments of adenylate cyclase. Fusion proteins were expressed from low-copy (pKNT25/pKT25) or high-copy (pUT18/pUT18C) plasmids as N-terminal (pKNT25/pUT18) or C-terminal (pKT25/pUT18C) of T18 or T25 fragments. Co-transformation of T18 and T25 fusions into *E. coli* BTH101 was done using the heat-shock procedure of calcium-chloride treated cells. Interactions were scored on LB plates supplemented with antibiotics (ampicillin and kanamycin), IPTG and X-Gal (5-bromo-4-chloro-3-indolyl-beta-D-galacto-pyranoside). As negative control, empty plasmid pK(N)T25 and pUT18(C) were also co-transformed into *E. coli* BTH101.

For checking the interaction of FlaA/B and Maf2 in tripartite system expressing GlfM, the GlfM coding sequence was placed on a compatible plasmid (pSRKGm-3xFlag-GlfM) or immediately downstream of T18/T25-Maf2 fusions.

Primers used for the cloning of the different ORFs as well as the resulting constructs are listed in **Supplementary Table S2** and **Supplementary Table S3**, respectively.

### Strain and plasmid construction

Plasmid recombinants were obtained by electroporation in *C. crescentus,* while for *S. oneidensis* plasmids were conjugated by bi-parental mating from the *E. coli* WM3064 donor cells. To construct in-frame deletion mutants, pNPTS138-R6KT^53^ derivatives constructed as detailed below were mobilized into *S. oneidensis* by bi-parental mating. Double recombinants were first selected on LB plates containing sucrose and then checked for sensitivity to kanamycin. Successful deletion was further checked in candidate clones by Sanger sequencing of the modified locus amplified by PCR using primers external to the homology DNA sequences used for recombination on pNPTS138-R6KT-derived plasmids (**Supplementary Table S2**).

pNPTSR6KT-Δ*maf1*: PCR was used to amplify two homology DNA fragments flanking the *maf1* ORF, by using primers So3273_T1F/So3273_T1R and So3273_T2F/So3273_T2R. PCR fragments were digested with *Spe*I/*EcoR*I and *EcoR*I/*Sal*I, respectively. Ultimately, fragments were ligated onto *Spe*I/*Sal*I-digested pNPTS138-R6KT.

pNPTSR6KT-Δ*maf2*: PCR was used to amplify two homology DNA fragments flanking the *maf2* ORF (So3259), by using primers So3259_T1F/So3259_T1R and So3259_T2F/So3259_T2R. PCR fragments were digested with *Spe*I/*EcoR*I and *EcoR*I/*Sal*I, respectively. Ultimately, fragments were ligated onto *Spe*I/*Sal*I-digested pNPTS138-R6KT.

pNPTSR6KT- Δ*pseI*: PCR was used to amplify two homology DNA fragments flanking the *pseI* ORF (So3261), by using primers T1-*Spe*I-So-*neuB*/T1-*EcoR*I-So-*neuB* and T2-*EcoR*I- So-*neuB*/T2-*Sal*I-So-*neuB*. PCR fragments were digested with *Spe*I/*EcoR*I and *EcoR*I/*Sal*I, respectively. Ultimately, fragments were ligated onto *Spe*I/*Sal*I-digested pNPTS138-R6KT.

pNPTSR6KT- Δ*pseC*: PCR was used to amplify two homology DNA fragments flanking the *pseC* ORF, by using primers T1-*Spe*I-*pseC*-So/T1-*EcoR*I-*pseC*-So and T2-*EcoR*I-pseC- So/T2-*Sal*I-*pseC*-So. PCR fragments were digested with *Spe*I/*EcoR*I and *EcoR*I/*Sal*I, respectively. Ultimately, fragments were ligated onto *Spe*I/*Sal*I-digested pNPTS138-R6KT.

pNPTSR6KT- Δ*flaA*: PCR was used to amplify two homology DNA fragments flanking the *flaA* ORF, by using primers T1-*Spe*I-So-*flaA*-F/T1-*Bam*HI-So-*flaA*-R and T2-*Bam*HI-So-*flaA*-F/T2- *Sal*I-So-*flaA*-R. PCR fragments were digested with *Spe*I/BamHI and BamHI/*Sal*I, respectively. Ultimately, fragments were ligated onto *Spe*I/*Sal*I-digested pNPTS138-R6KT.

pNPTSR6KT- Δ*glfM*: PCR was used to amplify two homology DNA fragments flanking the *glfM* ORF (So3260), by using primers *Spe*I-T1-So3260/*Bam*HI-T1-So3260 and *Bam*HI-T2- So3260/*Sal*I-T2-So3260. PCR fragments were digested with *Spe*I/*Bam*HI and *Bam*HI/*Sal*I, respectively. Ultimately, fragments were ligated onto *Spe*I/*Sal*I-digested pNPTS138-R6KT.

Expression, complementation and BACTH plasmids were constructed either by conventional restriction cloning, inverse PCR or Gibson assembly^54^ as detailed below. For inverse PCR, amplicons were subjected to 5’ end phosphorylation using T4 polynucleotide kinase (Thermofischer) prior to ligation using T4 DNA Ligase (Thermofischer). For Gibson assembly, DNA fragments were assembled using Nebuilder® Hifi DNA Assembly mix (New England Biolabs) following manufacturer’s instructions and incubation at 50°C for 75min.

pJU0001: The synthetic fragment encoding FlaA and Maf1 of *S. oneidensis*, codon-optimized for *E. coli*, was subcloned as *Nde*I/*Xba*I fragment onto pSRK-Gm.

pJU0002: The synthetic fragment encoding FlaB and Maf2 of *S. oneidensis*, codon-optimized for *E. coli*, was subcloned as *Nde*I/*Xba*I fragment onto pSRK-Gm.

pJU0005: The synthetic fragment encoding FliC and Maf of *Pseudomonas* sp. Irchel 3E13, codon-optimized for *E. coli*, was subcloned as *Nde*I/*Xba*I fragment onto pSRK-Gm.

pJU0006: To generate a pSRK-Gm expressing only a 3xFLAG-FlaA, *maf1* was removed from pJU001 by *Spe*I digestion followed by re-ligation of pJU0001.

pJU0007: To generate a pSRK-Gm expressing only a 3xFLAG-FlaB, *maf2* was removed from pJU0002 by *Spe*I digestion followed by re-ligation of pJU002.

pJU0008: To add an N-terminal HA-tag in Maf1, inverse PCR using primers HA-So-Maf1syn- For/So-FlaAsyn-Rev was done on pJU0006.

pJU0009: To add an N-terminal HA-tag in Maf2, inverse PCR using primers HA-Maf2syn- For/So-FlaBsyn-Rev was done on pJU0002.

pJU0010: To insert HA-Maf1 downstream of 3xFLAG-*flaB* on pJU007, HA-Maf1 was excised from pJU0008 and subcloned as *Spe*I/*Pst*I fragment onto pJU0007.

pJU0011: To introduce the D247A point mutation in *maf1*, inverse PCR using So3273-D247A- For/So3273-D247A-Rev was done on pJU0001.

pJU0012: To introduce the D333A point mutation in *maf1*, inverse PCR using So3273-D333A- For/So3273-D333A-Rev was done on pJU0001.

pJU0013: To add an N-terminal HA-tag in *maf1*-D247A, inverse PCR using primers HA-So- Maf1syn-For/So-FlaAsyn-Rev was done on pJU0011.

pJU0014: To add an N-terminal HA-tag in *maf1*-D333A, inverse PCR using primers HA-So- Maf1syn-For/So-FlaAsyn-Rev was done on pJU0012.

pJU0015: To introduce the D261A point mutation in *maf2*, inverse PCR using SoMaf2syn- D261A-For/SoMaf2syn-D261A-Rev was done on pJU0009.

pJU0016: To introduce the D347A point mutation in *maf2*, inverse PCR using SoMaf2syn- D347A-For/SoMaf2syn-D347A-Rev was done on pJU0009.

pJU0017, pJU0018 and pJU0019: To delete 3xFLAG-FlaB, inverse PCR was done using primers pSRKGm-INVRS1/pSRKGm-INVRS2 on pJU0009, pJU0015 and pJU0016, respectively.

pJU0020: The truncated *maf1* (1-420 aa) of *S. oneidensis* was generated by PCR using primers So-FlaAsyn-int-For/PstI-So3273-1-420-Rev from pJU008. The subsequent amplicon was cloned as *Spe*I-*Pst*I fragment onto pJU0008 replacing the full-length HA-Maf1.

pJU0021: The truncated *maf1* (1-420 aa) of *S. oneidensis* was cloned onto pJU0007 as *Spe*I-*Pst*I fragment.

pJU0022: Inverse PCR was done to generate a truncated *maf1* (1-699 aa) using primers pSRK-HA-Maf1-1-699aa-INVRS1/pSRK-HA-Maf1-1-699aa-INVRS2 and pJU0008 as template.

pJU0023: Inverse PCR was done to generate a truncated *maf1* (1-699 aa) using primers pSRK-HA-Maf1-1-699aa-INVRS1/pSRK-HA-Maf1-1-699aa-INVRS2 and pJU0010 as template.

pJU0024: HA-Maf2 was amplified from pJU0009 using So-Flg-int-Rev/SalI-HA-SoMaf2 and cloned as *Spe*I-*Sal*I fragment onto pJU0006.

pJU0025: HA-Maf2 was amplified from pJU0009 using So-Flg-int-Rev/SalI-SoMaf2 and cloned as *Spe*I-*Sal*I fragment onto pJU0008.

pJU0026: A synthetic fragment encoding 3xFLAG-FlaA which harbours alanine substitutions in five serine residues (S105, S143, S171, S180 and S185) and subcloned as *Nde*I-*Spe*I fragment onto pSRK-Gm.

pJU0027: A synthetic fragment encoding 3xFLAG-FlaA which harbours alanine substitutions in two serine residues (S105 and S180) subcloned as *Nde*I-*Xba*I fragment onto pSRK-Gm.

pJU0028: A synthetic fragment encoding3xFLAG-FlaA which harbours alanine substituions in three serine residues (S105, SI71 and S180) subcloned as *Nde*I-*Xba*I fragment onto pSRK-Gm.

pJU0029: A synthetic fragment encoding3xFLAG-FlaA which harbours alanine substitutions in two serine residues (S143 and S185) subcloned as *Nde*I-*Xba*I fragment onto pSRK-Gm.

pJU0030: A synthetic fragment encoding 3xFLAG-FlaA which harbours alanine substitutions in three serine residues (S143, SI71 and S185) subcloned as *Nde*I-*Xba*I fragment onto pSRK- Gm.

pJU0031: CDS of *glfM* (So3260) was assembled onto pSRK-Gm by Gibson assembly. CDS of *glfM* was amplified from *S. oneidensis* genome using primers GB-pSRK-3xFLSo3260-1/GB- pSRK-3xFLSo3260-2 while pSRK-Gm harbouring a 3xFLAG was amplified from pJU0006 using primers GB-pSRK-3xFLSo3260-3/GB-pSRK-3xFLSo3260-4.

pJU0032: CDS of 3xFLAG-*glfM* was amplified from pJU00031 using primers SRK-For/SRK- Rev and subcloned as a *Nde*I-*Kpn*I fragment onto pMT375.

pJU0033: A synthetic fragment encoding *S. oneidensis glfM* (GlfM-F24L-SPOT) harbouring a F24L mutation and a C-terminal SPOT-tag, codon-optimized for *E. coli* and subcloned as a *Nde*I/*Xba*I fragment onto pSRK-Gm.

pJU0034: A synthetic fragment encoding *S. oneidensis glfM* (GlfM-R25K-SPOT) harbouring a R25K mutation and a C-terminal SPOT-tag, codon-optimized for E. coli and subcloned as a *Nde*I/*Xba*I fragment onto pSRK-Gm.

pJU0035: A synthetic fragment encoding *S. oneidensis glfM* (GlfM-D77A-SPOT) harbouring a D77A mutation and a C-terminal SPOT-tag, codon-optimized for *E. coli* and subcloned as a *Nde*I/*Xba*I fragment onto pSRK-Gm.

pJU0036: A synthetic fragment encoding *S. oneidensis glfM* (GlfM-E80A-SPOT) harbouring a E80A mutation and a C-terminal SPOT-tag, codon-optimized for *E. coli* and subcloned as a *Nde*I/*Xba*I fragment onto pSRK-Gm.

pJU0037: The D77A point mutation was introduced into 3xFLAG-GlfM carried by pJU0028 by inverse PCR using primers pSRK-3xFL-So3260-INVRS1/pSRK-3xFL-So3260-INVRS2.

pJU0038 and pJU0039: CDS of *glfM* (CKW37_RS17405) from *Pseudomonas sp*. Irchel 3E13 (GenBank: NZ_FYDX01000009.1) was amplified from its genome using primers NdeI-3E13- fliSlike-For/KpnI-3E13-fliSlike-Rev and cloned as a *Nde*I/*Kpn*I fragment onto pSRK-Gm and pMT375, respectively.

pJU0040 and pJU0041: CDS of *glfM* (SPUTCN32_RS13550) from *Shewanella putrefaciens* CN-32 (GenBank: NC_009438.1) was amplified from its genome using primers NdeI-SpCN32- GlfM-For/KpnI-SpCN32-GlfM-Rev and cloned as a *Nde*I/*Kpn*I fragment onto pSRK-Gm and pMT375, respectively.

pJU0042 and pJU0043: CDS of *glfM* (NCTC10735_01035) from *Shewanella baltica* NCTC10735 (GenBank: NZ_UGYM01000002.1) was amplified from its genome using primers NdeI-Sb-GlfM/KpnI-Sb-GlfM and cloned as a *Nde*I/*Kpn*I fragment onto pSRK-Gm and pMT375, respectively.

pJU0044 and pJU0045: CDS of *glfM* from *Kurthia* sp. 11kri321 (GenBank: NZ_CP013217.1), codon-optimized for *E. coli*, was amplified from a synthetic fragment *xyl*-Ku-Maf-GlfM-Hag- HAH6 using primers NdeI-Ku-GlfM/KpnI-Ku-GlfM and subcloned as *Nde*I/*Kpn*I fragment onto pSRK-Gm and pMT375, respectively.

pJU0046: A synthetic fragment (Ku-Maf-GlfM-HagHAH6) encoding the Maf, GlfM and Hag flagellin of *Kurthia* sp. 11kri321, codon-optimized for *E. coli* and subcloned as *Nde*I/*Xba*I fragment onto pSRK-Gm.

pJU0047: *glfM* of *Kurthia* was removed from pJU00046 by inverse PCR using primers Ku-Maf- INVRS1/Ku-Hag-INVRS2.

pJU0048 and pJU0049: CDS of FlaA (sans START and STOP) was amplified from *S. oneidensis* using pUT-pK-XbaI-FlaA-F/pUT-pK-BamHI-FlaA-R and cloned respectively as a *Xba*I-*Bam*HI fragment onto pUT18 and pUT18C, in-framed of the T18 fragment.

pJU0050 and pJU0051: CDS of FlaA (sans START and STOP) was amplified from *S. oneidensis* using pUT-pK-XbaI-FlaA-F/pUT-pK-BamHI-FlaA-R and cloned respectively as a *Xba*I-*Bam*HI fragment onto pKNT25 and pKT25, in-framed of the T25 fragment.

pJU0052 and pJU0053: CDS of FlaB (sans START and STOP) was amplified from *S. oneidensis* using pUT-pK-XbaI-FlaB-F/pUT-pK-BamHI-FlaB-R and cloned rescpectively as a *Xba*I-*Bam*HI fragment onto pUT18 and pUT18C, in-framed of the T18 fragment.

pJU0054 and pJU0055: CDS of FlaB (sans START and STOP) was amplified from *S. oneidensis* using pUT-pK-XbaI-FlaB-F/pUT-pK-BamHI-FlaB-R and cloned respectively as a *Xba*I-*Bam*HI fragment onto pKNT25 and pKT25, in-framed of the T25 fragment.

pJU0056 and pJU0057: CDS of Maf1 (sans START and STOP) was amplified from *S. oneidensis* using pKNT25-pUT-PstI-Maf1-F/pK-pUT-KpnI-Maf1-R and cloned respectively as a *Pst*I-*Kpn*I fragment onto pUT18 and pUT18C, in-framed of the T18 fragment.

pJU0058: CDS of Maf1 (sans START and STOP) was amplified from *S. oneidensis* using primers pKNT25-pUT-PstI-Maf1-F/pK-pUT-KpnI-Maf1-R and cloned as a *Pst*I-*Kpn*I fragment onto pKNT25, in-framed of the T25 fragment.

pJU0059: CDS of Maf1 (sans START and STOP) was amplified from *S. oneidensis* using primers pKT-PstI-Maf1-F/pK-pUT-KpnI-Maf1-R and cloned as a *Pst*I-*Kpn*I fragment onto pKT25, in-framed of the T25 fragment.

pJU0060 and pJU0061: CDS of Maf2 (sans START and STOP) was amplified from *S. oneidensis* using primers pKNT25-pUT-PstI-Maf2-F/pK-pUT-KpnI-Maf2-R and cloned respectively as a *Pst*I-*Kpn*I fragment onto pUT18 and pUT18C, in-framed of the T18 fragment.

pJU0062: CDS of Maf2 (sans START and STOP) was amplified from *S. oneidensis* using primers pKNT25-pUT-PstI-Maf2-F/pK-pUT-KpnI-Maf2-R and cloned as a *Pst*I-*Kpn*I fragment onto pKNT25, in-framed of the T25 fragment.

pJU0063: CDS of Maf2 (sans START and STOP) was amplified from *S. oneidensis* using primers pKT25-PstI-Maf2-F/pK-pUT-KpnI-Maf2-R and cloned as *Pst*I-*Kpn*I fragment onto pKT25, in-framed of the T25 fragment.

pJU0064 and pJU0065: Truncated Maf1 (1-420 aa) was amplified from *S. oneidensis* using primers PstI-SoMaf1-F/pK-pUT-KpnI-maf1trunc-Rev and cloned respectively as a *Pst*I-*Kpn*I fragment onto pUT18 and pUT18C, in-framed of the T18 fragment.

pJU0066: Truncated Maf1 (1-420 aa) was amplified from *S. oneidensis* using primers PstI- SoMaf1-F/pK-pUT-KpnI-maf1trunc-Rev and cloned as a *Pst*I-*Kpn*I fragment onto pKNT25, in- framed of the T25 fragment.

pJU0067: Truncated Maf1 (1-420 aa) was amplified using pKT25-PstI-maf1-F/pK-pUT-KpnI- maf1trunc-Rev and cloned as a *Pst*I-*Kpn*I fragment onto pKT25, in-framed of T25 fragment.

pJU0068 and pJU0069: CTD (421-822 aa) of Maf1 was amplified from *S. oneidensis* using primers pUT-pKN-PstI-CTD2wt-For/pK-pUT-KpnI-maf1CTD2wt-Rev and cloned respectively as a *Pst*I-*Kpn*I fragment onto pUT18 and pUT18C, in-framed of the T18 fragment.

pJU0070: CTD (421-822 aa) of Maf1 was amplified from *S. oneidensis* using primers pUT- pKN-PstI-CTD2wt-For/pK-pUT-KpnI-maf1CTD2wt-Rev and cloned as a *Pst*I-*Kpn*I fragment onto pKNT25, in-framed of the T25 fragment.

pJU0071: CTD (421-822 aa) of Maf1 was amplified from *S. oneidensis* using pKT- Maf1CTD2wt-For/pK-pUT-KpnI-maf1CTD2wt-Rev and cloned as a PstI-KpnI fragment onto pKT25, in-framed of T25 fragment.

pJU0072 and pJU0073: CDS of GlfM (sans START and STOP) was amplified from *S. oneidensis* using primers XbaI-So3260-For/BamHI-So3260-Rev and cloned respectively as a *Xba*I-*Bam*HI fragment onto pUT18 and pUT18C, in-framed of T18 fragment.

pJU0074 and pJU0075: CDS of GlfM (sans START and STOP) was amplified from *S. oneidensis* using primers XbaI-So3260-For/BamHI-So3260-Rev and cloned respectively as a *Xba*I-*Bam*HI fragment onto pKNT25 and pKT25, in-framed of T25 fragment.

pJU0076: CDS of GlfM (full-length) was assembled onto pJU0060 by Gibson assembly. The CDS was amplified from *S. oneidensis* using primers GB-So3260-T18-For/GB-So3260- T18CMaf2-Rev while pJU0060 was amplified using GB-T18CMaf2-So3260-For/GB-T18- So3260-Rev.

pJU0077: CDS of GlfM (full-length) was assembled onto pJU0061 by Gibson assembly. The CDS was amplified from *S. oneidensis* using primers GB-So3260-T18-For/GB-So3260- T18CMaf2-Rev while pJU0061 was amplified using GB-T18CMaf2-Fo/GB-T18CMaf2- So3260-For.

pJU0078: CDS of GlfM (full-length) was assembled onto pJU0062 by Gibson assembly. The CDS was amplified from *S. oneidensis* using primers GB-So3260-pKN-For/GB-So3260- T18CMaf2-Rev while pJU0062 was amplified using GB-T18CMaf2-So3260-F/GB-pKN- So3260-Rev.

pJU0079: CDS of GlfM (full-length) was assembled onto pJU0063 by Gibson assembly. The CDS was amplified from *S. oneidensis* using primers GB-So3260-pKT-For/GB-So3260-pKT- Rev while pJU0063 was amplified using GB-pKT-So3260-For/GB-pKT-So3260-Rev.

All plasmids, synthetic fragments and primers are listed in **Appendix Table 1, Appendix Table 2 and Appendix Table 3**, respectively.

## SUPPLEMENTAR FIGURE LEGENDS

**Supplementary Figure S1.**
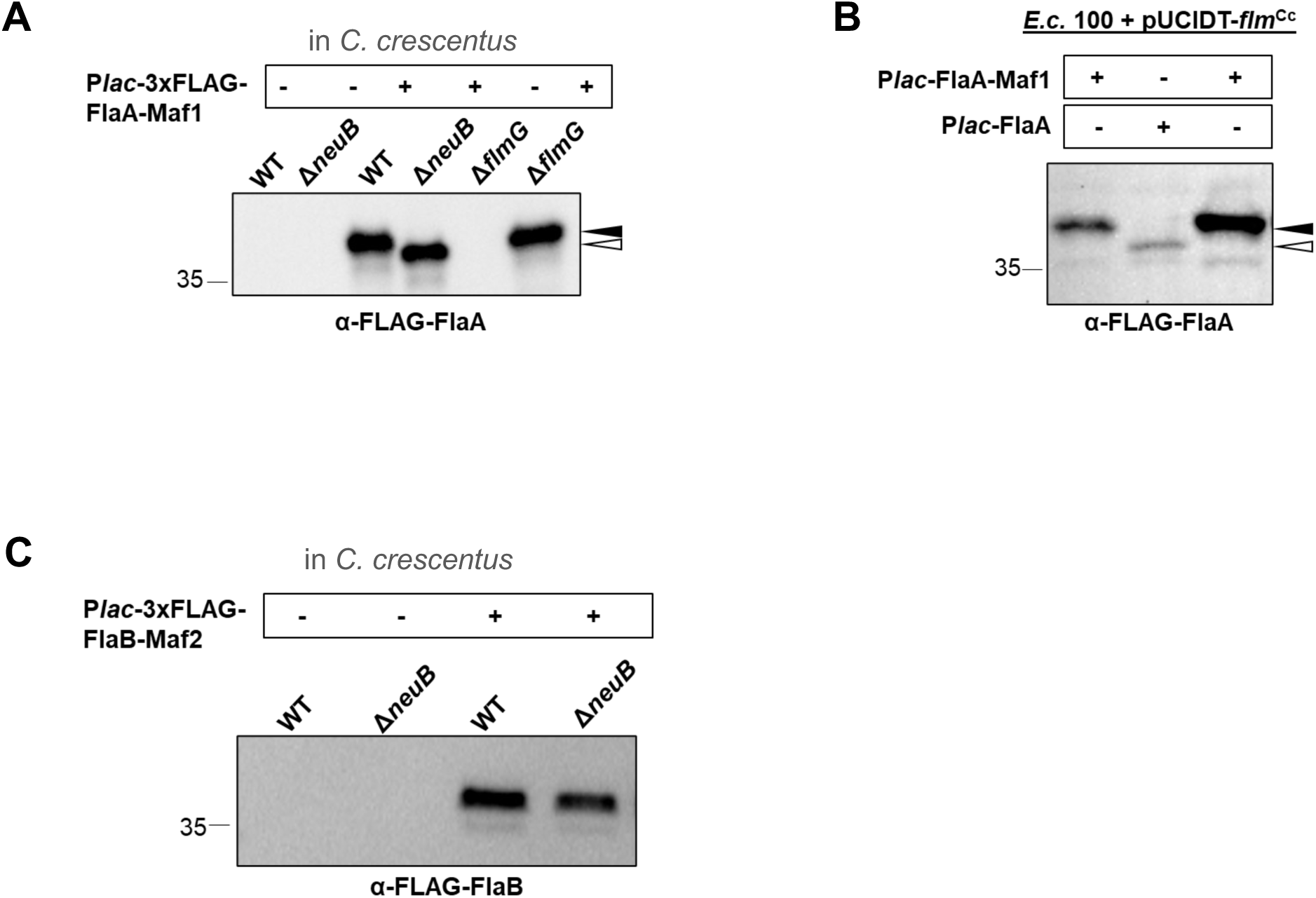
Reconstitution of flagellin glycosylation by Maf-1 and Maf-2 in heterologous hosts. **A)** Immunoblot of 3xFLAG-tagged FlaA of *S. oneidensis* showing Maf-1 and Pse- dependent migration changes of FLAG-FlaA in *C. crescentus WT* or Δ*flmG* that produce Pse *versus* Δ*neuB* cells that lack Pse. Blots were probed with monoclonal antibodies to the FLAG-tag (α-FLAG-FlaA). **B)** Immunoblot of 3xFLAG-tagged FlaA of *S. oneidensis* showing reconstitution of FLAG- FlaA glycosylation by Maf-1 in the *E. coli* cloning strain EC100D producing the synthetic *flm* operon of *C. crescentus*. The change in FLAG-FlaA migration was revealed by probing blots with monoclonal antibodies to the FLAG-tag (α-FLAG-FlaA). **C)** Immunoblot of 3xFLAG-tagged FlaB of *S. oneidensis* showing that FLAG-FlaB does not change its migration in *C. crescentus WT* or Δ*neuB* cells co-expressing Maf-2. Pse is produced in *WT* cells but not in Δ*neuB* cells.

**Supplementary Figure S2.**
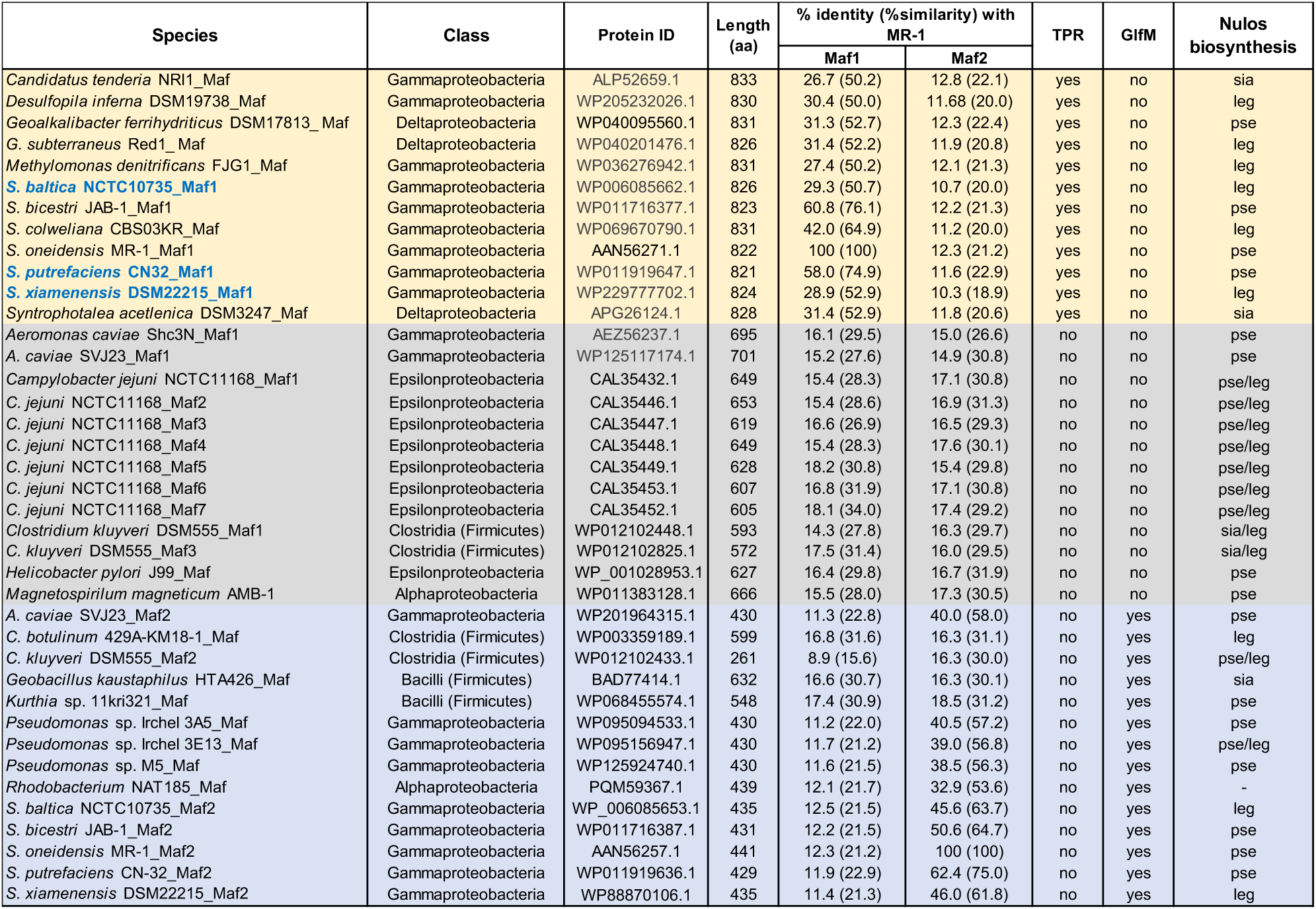
Pairwise amino acid sequence comparison between Maf- 1/Maf-2 of *S. oneidensis* MR-1 and other existing Maf types. Global sequence alignment was done using EMBOSS Needle at EMBL-EBI. For sequences sharing below 35% identity to MR-1 Mafs, BLOSUM35 was used as substitution matrix, while BLOSUM60 for above 35%. Different Maf types were categorized based on the presence of a TPR domain (light yellow), absence of a TPR domain (light grey) and presence of GlfM (light blue). Species that are in bold blue font were used for the complementation of Δ*maf1*.

**Supplementary Figure S3.**
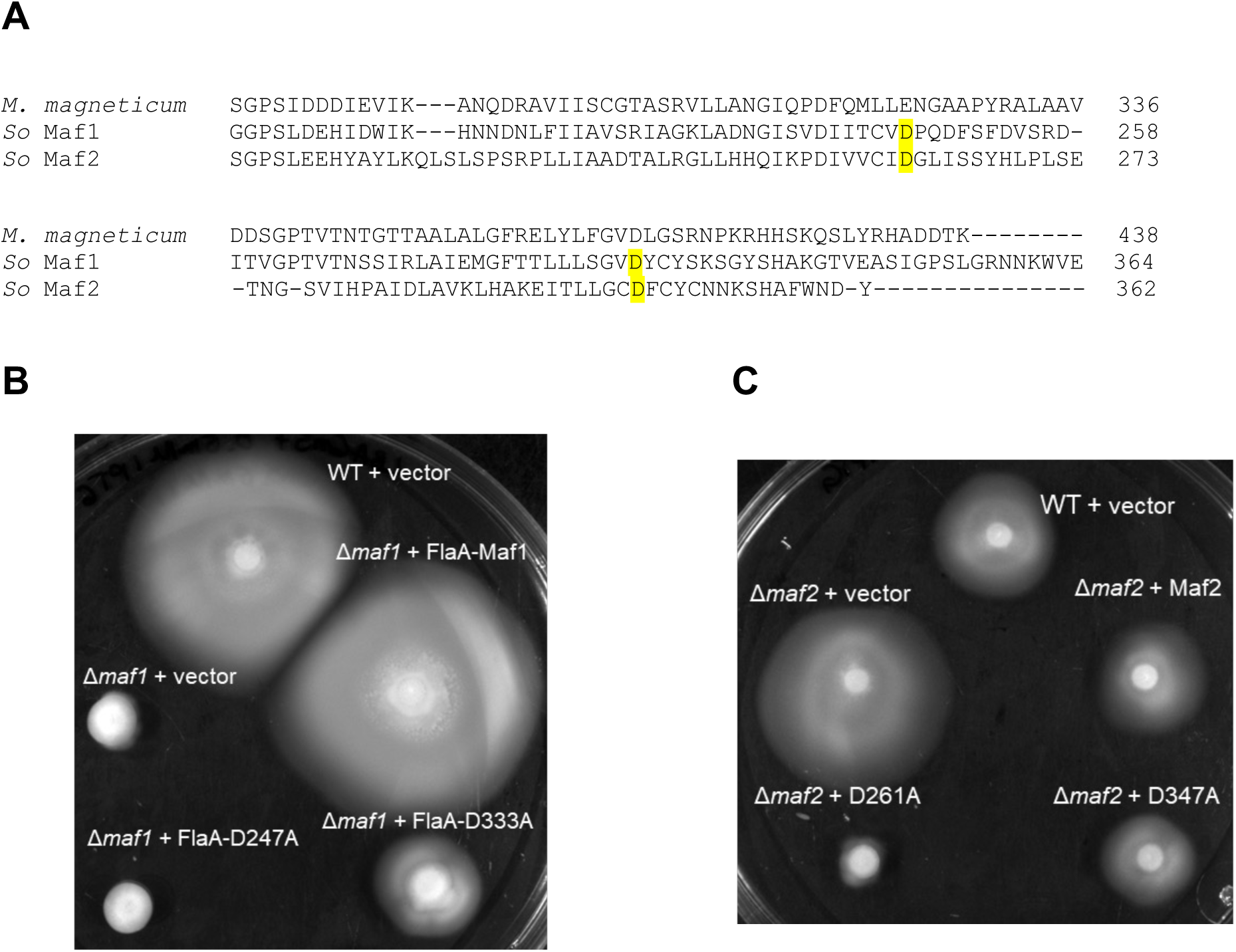
Essential residues involved in the catalytic activity of Maf-1 and Maf-2. **A)** Sequence alignment of *S. oneidensis* Maf-1 and Maf-2 with *Magnetospirilum magneticum* AMB-1 Maf using MUSCLE at EMBL-EBI. Amino acid residues substituted with alanine are highlighted in yellow. **B)** Motility assay of Δ*maf1* mutant complemented with point mutants of Maf1 on LB swarm (0.25%) agar. **C)** Motility assay of Δ*maf2* mutant complemented with point mutants of Maf2 on LB swarm (0.25%) agar.

**Supplementary Figure S4.**
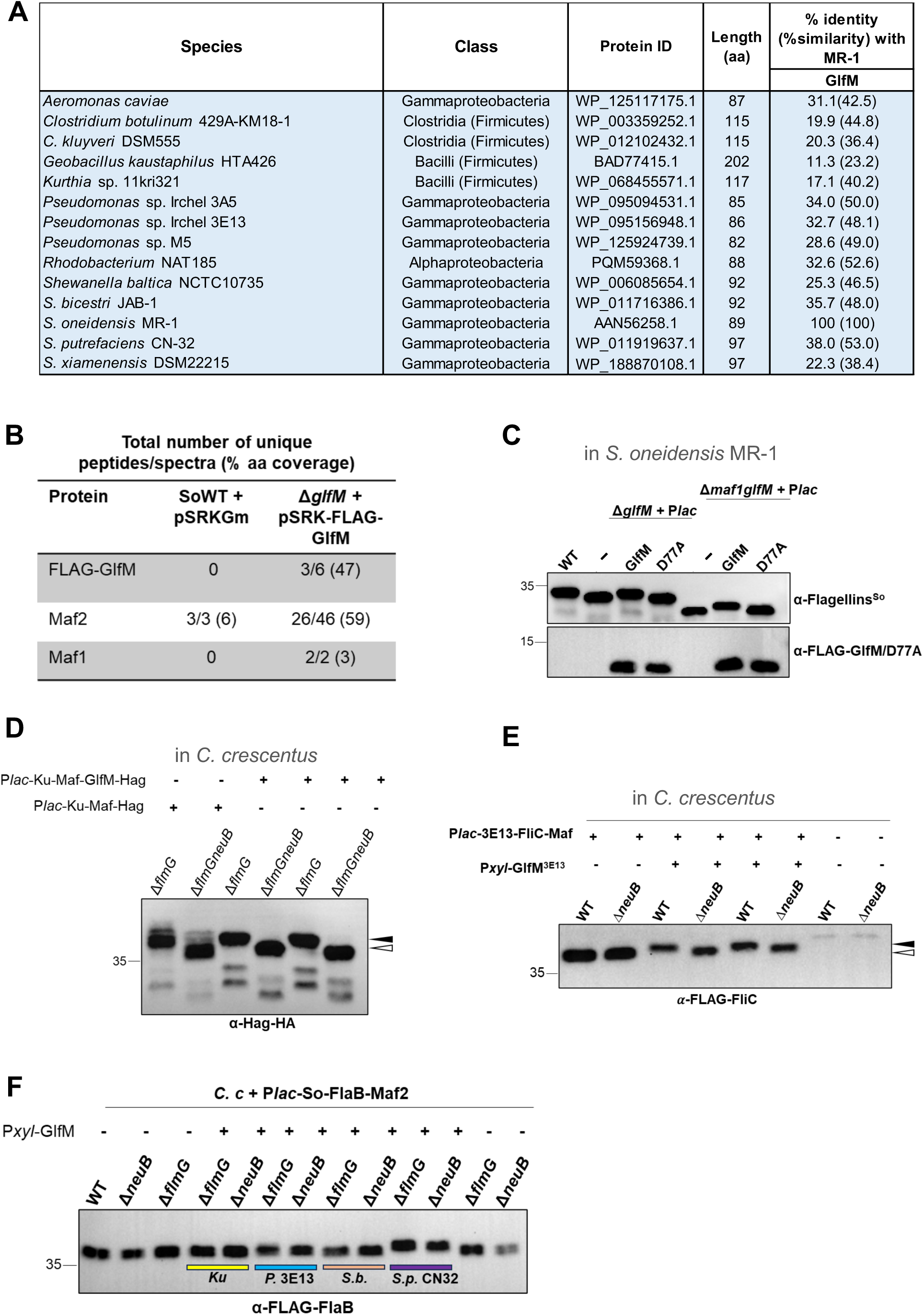
Similarity, co-immunoprecipitation and functional conservation of GlfM. **A)** Pairwise amino acid sequence alignment between GlfM of MR-1 and orthologs. Global sequence alignment was done using EMBOSS Needle algorithm at EMBL-EBI. For sequences sharing below 35% identity to MR-1 GlfM, BLOSUM35 was used as substitution matrix, while BLOSUM60 for above 35%. **B)** Mass spectrometry (LC-MS/MS) analysis of 3xFLAG-tagged GlfM (FLAG-GlfM) pulled-down from extracts of complemented Δ*glfM* cells. As negative control, *S. oneidensis* WT expressing the empty vector pSRK-Gm was used. Data shown are the total number of unique peptides and spectra as wells % amino acid coverage of the bait FLAG-GlfM and the detected Maf-2 and Maf-1 based on the following criteria in Scaffold: protein threshold at 1% FDR, peptide threshold at 0.1% FDR and minimum of 2 peptides. FDR corresponds to False Discovery Rate. **C)** Immunoblot of *S. oneidensis* flagellins as well as of the 3xFLAG-tagged GlfM (FLAG- GlfM) and the point mutant GlfM-D77A (FLAG-GlfM-D77A) expressed in Δ*glfM* single mutant and Δ*maf-1*Δ*glfM* double mutant cells. Blots were with monoclonal antibodies to the FLAG-tag (α-FLAG-GlfM /D77A) and with polyclonal antibodies to *S. oneidensis* FlaA/B (α-Flagellins^So^). **D)** Immunoblot of HA-tagged Hag (Hag-HA) flagellin of *Kurthia* sp. 11kr321 expressed in *C. crescentus* showing a Maf- and Pse-dependent reduction in migration of *Kurthia* Hag-HA even in the absence of GlfM, but GlfM promotes this change in migration. Blots were probed with monoclonal antibodies to the HA-tag (α-Hag-HA). White carets indicate the migration of unmodified Hag-HA, whereas black carets indicate the slower migration of modified Hag-HA. **E)** Immunoblot of 3xFLAG-tagged FliC flagellin (FLAG-FliC) of *Pseudomonas* sp. 3E13 showing a GlfM-Maf-2 modification of FLAG-FliC in *C. crescentus WT* cells that produce Pse versus Δ*neuB* cells that lack Pse. Note that expression of GlfM from *Pseudomonas* sp. 3E13 (GlfM^3E13^) results in a Pse-dependent reduction in migration of FLAG-FliC as indicated by the shift from the white to the black carets, respectively. Blots were probed with monoclonal antibodies to the FLAG-tag (α-FLAG-FliC). **F)** Immunoblot of 3xFLAG-tagged FlaB (FLAG-FlaB) from *S. oneidensis* in *C. crescentus* cells co-expressing GlfM othologs from *Kurthia* sp. 11kr321 (*Ku*), *Pseudomonas* sp. 3E13 (*P.3E13*), *Shewanella baltica* (*S. b.*) and *Shewanella putrefaciens* (*S. p.* CN32). Blots were probed with monoclonal antibodies to the FLAG-tag (α-FLAG-FlaB).

